# High-throughput stomatal phenotyping provides selection targets for stress-resilient wheat

**DOI:** 10.64898/2026.07.10.737162

**Authors:** Mahmoud Mabrouk, Nicholas J. Russell, Emilio Villar Alegria, Tien-Cheng Wang, Jui-An Liang, Fang-Jin Wu, Yunfeng Huang, Benjamin Wittkop, Rod Snowdon, Lukas Förter, Anna Moritz, Eva Herzog, Eliyeh Ganji, Gwendolin Wehner, Andreas Stahl, Tsu-Wei Chen

## Abstract

Phenotyping stomatal traits and their developmental plasticity is time-consuming but holds potential to improve water use efficiency and photosynthesis for designing stress-tolerant crops under climate change. Here, we develop a robust, high-throughput pipeline for phenotyping 14 stomatal traits in winter wheat related to size, variation, maximum conductance, and spatial patterning. We (1) analyze over 25,000 images from 60 wheat cultivars grown in growth chamber, greenhouse, and field conditions; (2) investigate the impact of light, temperature, and reduced water and nitrogen supply on stomatal traits and their developmental plasticity across adaxial and abaxial surfaces; and (3) evaluate genetic diversity and breeding progress of stomatal traits. Stomatal traits were highly broad-sense heritable, were largely plastic in response to environmental conditions, and showed genotype-specific responses. Stomatal traits of third leaves under controlled environments with stable light and temperature conditions reliably captured the genetic variance of flag leaves under field conditions. Our data suggests that the upper leaf surface contributed more to transpiration and cooling through consistently higher stomatal density, area, and maximum conductance, while the lower surface facilitated CO₂ diffusion via systematic proper patterning and spacing. Breeding maintains the genetic diversity of stomatal traits, and our pipeline facilitates breeders to target them to enhance water use efficiency in high-yielding modern cultivars.

## Introduction

Stomatal traits play a substantial role in regulating the trade-off between CO₂ uptake for photosynthesis and water loss via transpiration. They maintain the balance between primary productivity and water-use efficiency (WUE) across agricultural and natural ecosystems.^1^ Under conditions of limited water availability or elevated demand for photo-assimilates, such as during periods of high temperature in rapid growing periods, smart strategies to control gas exchange become critical to sustaining productivity while mitigating drought stress.^2,3^

Stomatal morphology plays a critical role in stomatal regulation. Stomatal geometry (depth, length, and width) significantly influences gas exchange and photosynthetic efficiency. For example, stomatal conductance is fundamentally regulated by both anatomical characteristics, such as stomatal density, size (guard cell length), and patterning, and functional responses that regulate pore aperture in response to environmental stimuli, such as light and water stress.^4–10^ Moreover, a more favorable width-to-length ratio can reduce resistance to gas diffusion, thereby enhancing conductance.^11^ Beyond individual geometry, stomatal patterning (i.e., the spatial arrangement of stomata) further affects stomatal function. Guard cells depend on ionic exchange with neighboring epidermal cells to regulate aperture dynamics, enabling effective stomatal opening and closing.^12–14^ Disruption in this spatial relationship can impair function. For instance, genotypes exhibiting closely spaced and highly clustered stomata consistently show reduced maximum stomatal conductance (*g*_smax_) compared to those with uniformly distributed stomata of similar density.^15^ This decline is likely due to reduced access to the surrounding epidermal cells, limiting ion exchange and compromising the mechanical flexibility required for efficient guard cell movement.^1,16^ Moreover, greater distances of stomata from their local center of gravity (referred to as stomatal divergence) may influence the efficiency of CO₂ diffusion across the epidermis and contribute to more effective leaf cooling by facilitating heat dissipation.^15^

Stomatal opening and closure are influenced by multiple factors, such as environmental triggers and morphological traits.^3,7,17,18^ Under saturating light, stomata open widely to meet photosynthetic demand. However, during a transition to low light, closure is initiated but proceeds more slowly, lagging behind the decline in photosynthesis and leading to unnecessary water loss.^7,13^ The speed of stomatal dynamics in response to environmental cues is not only shaped by biochemical processes (such as osmotic adjustments) but also by structural characteristics, including stomatal morphology and size.^1,19,20^ For example, dumbbell-shaped guard cells typically respond faster than elliptical ones, contributing to greater water-use efficiency.^13^ Similarly, smaller stomata generally exhibit faster dynamics, helping maintain gas exchange while improving long-term WUE.^19,21^ Interestingly, major cereal crops such as bread wheat and related species differ in their stomatal morphology and response speed to light.^10^ Stomatal regulation and WUE are strongly influenced by stomatal morphological traits (e.g., density and size), and the plasticity of a genotype to adjust them can shape its fitness under different environmental conditions.^22,23^ Therefore, understanding the genetic diversity of these traits and their plastic responses to different environmental triggers is essential.^20^

However, the extent of genetic diversity in stomatal traits among elite wheat cultivars remains largely unknown, mainly due to the time-consuming nature of such measurements. Traditionally, sample preparation to phenotype stomatal traits is labor-intensive.^24,25^ In recent years, stomatal phenotyping has advanced through non-destructive, high-throughput, high-resolution microscopy in combination with deep learning tools that enable efficient quantification of stomatal traits.^24,26–35^ Despite these advances, the genetic diversity of stomatal morphology in crop species remains poorly understood. Monocotyledonous leaves (e.g., wheat) have a linear blade with a V-shaped cross-section that rolls under stress due to shrinkage in bulliform cells^36,37^, preventing full leaf flattening and reducing image quality in high-throughput microscopy. This limits accurate quantification of traits such as stomatal characteristics^31^ and constrains the generation of high-quality phenotypic data, a key requirement for large-scale genotype selection and plant breeding improvement.^38,39^

As well, the number of high-resolution stomatal images at 400× magnification remains limited, particularly for wheat and species with small stomata or overlapping epidermal trichomes.^31,40^ Therefore, combining high-resolution imaging (e.g., 400×) with efficient computer vision and computational pipelines holds great potential for uncovering genetic diversity and heritability of stomatal traits, along with their plasticity in response to varying environmental conditions (light and temperature) or resource availability (water and nitrogen). Moreover, no systematic investigation has examined whether breeding has altered these traits, even though it has been suggested that in maize, stomatal conductance at the canopy level has not been a direct target of selection.^41^

In this study, we develop an accurate high-throughput method for phenotyping stomatal morphology in wheat, paired with an automated computational pipeline that quantifies 14 stomatal traits related to size, deviation, coverage, and spatial patterning. Using this pipeline, we investigate genetic diversity in stomatal morphology in sixty winter wheat cultivars in four growth chambers, three greenhouse experiments, and one field trial to i) assess the effects of light, temperature, drought stress, and nitrogen availability on stomatal morphological traits; ii) identify the plasticity of stomatal traits and how comparable they are under controlled conditions and a natural field environment; iii) explore the coordination in stomatal traits on abaxial and adaxial sides; and iv) determine which stomatal traits have been under-targeted by breeders’ selection using phenomics and genomics approaches. Our work underscores the importance and potential of high-throughput stomatal phenotyping to uncover subtle, yet crucial genotypic differences in breeding progress and leaf traits.

## Results

### An experimental pipeline assesses the effect of genotypic and environmental factors on stomatal morphology

We used sixty winter wheat cultivars (*Triticum aestivum* L.) to demonstrate the genotypic variation in responses to light and temperature. Fifty of the studied cultivars represent the breeding history of winter wheat in Germany between 1966 and 2018 (hereafter, the breeding panel), and ten represent exotic cultivars between 1963 and 1992 (hereafter, the exotic panel). These cultivars selected for the breeding panel were selected to adequately represent the breeding progress in Germany (see Methods and Table S1).^42^

We conducted experiments using several environmental conditions in growth chambers (GC), greenhouses (GH), and in the field (Fig. 1A). For GC experiments, we investigated all sixty genotypes in four different conditions: high temperature and high light intensity (HTHL), high temperature and low light intensity (HTLL), low temperature and high light intensity (LTHL), and low temperature and low light intensity (LTLH). Next, we examined the breeding panel in three greenhouse experiments under drought stress, constant temperature conditions, and 12-hour light-dark conditions. The first trial (GH1) consisted of thirty genotypes; the second trial (GH2) consisted of the remaining twenty genotypes and ten genotypes from the first trial chosen due to their varied water use efficiency (WUE, see Methods); and, the final trial (GH3) consisted of the ten genotypes which were tested in the previous two greenhouse experiments, and also examined their response to nitrogen availability along with drought stress (see Methods). Lastly, we conducted an open-field experiment for the breeding panel at the Groß-Gerau experimental station of Giessen University, Giessen, Germany (see Methods and Table S2).

**Figure 1:**
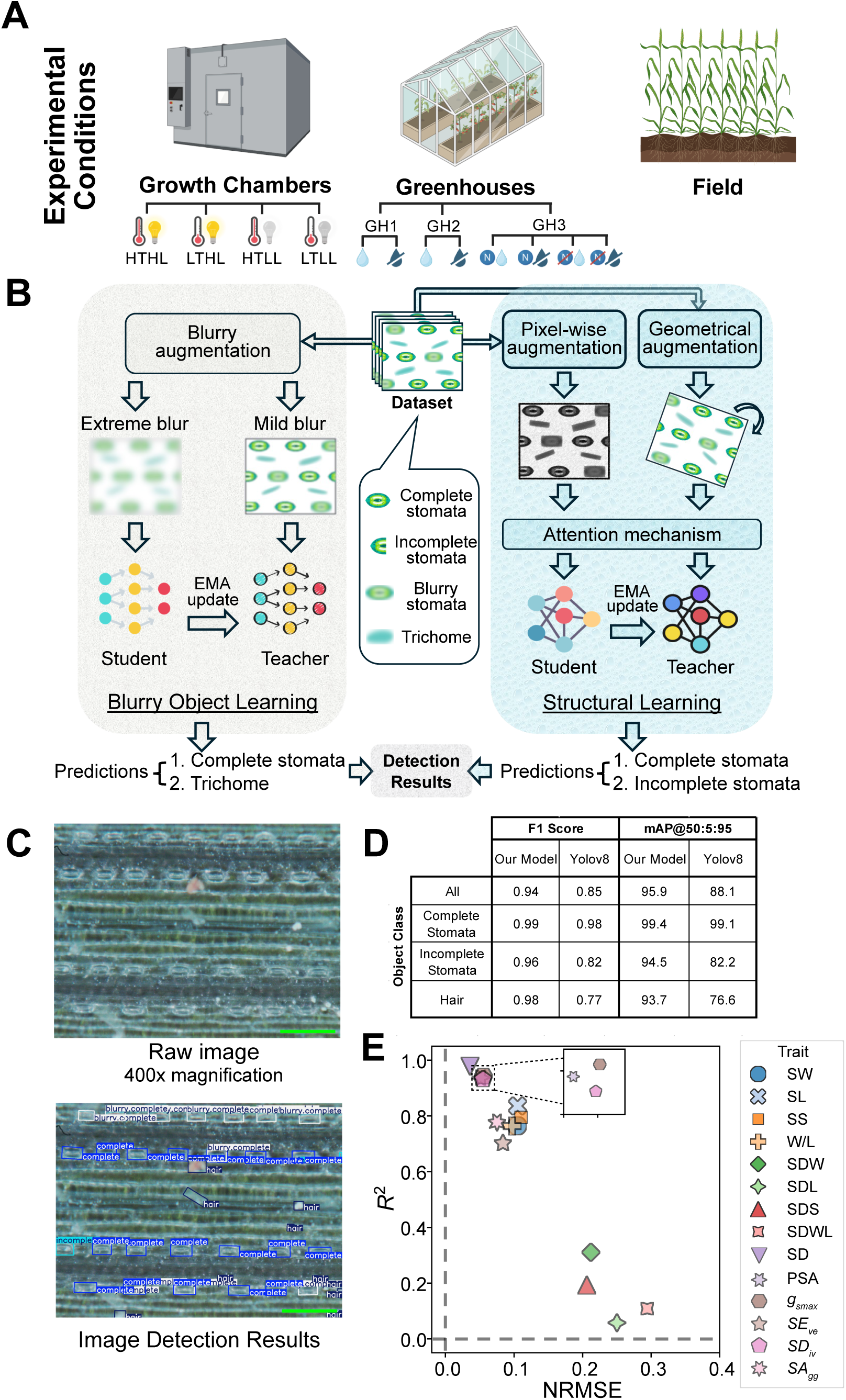
A high-throughput imaging pipeline accurately detects, classifies, and measures stomata from wheat. **(A)** An overview of the experimental setup. In brief, winter wheat was grown in growth chambers (GC), greenhouses (GH), and in the field. Light and temperature were varied in the GC experiments, while nitrogen and water availability were varied in the GH experiments. **(B)** An overview of the computational pipeline for detecting and classifying stomata from high-resolution images of wheat leaves. See Results and Methods sections for an in-depth explanation. In brief, using a manually labeled dataset of 504 images, the model was trained using a student-teacher framework using an attention mechanism and blurry object learning to find objects and classify them into one of the following: a complete stomata, an incomplete stomata, a blurry complete stomata, a blurry incomplete stomata, or a hair/trichome. (**C)** Above: representative microscope image of the abaxial side of a leaf; Below: identification of stomata and hair classes by the computational pipeline described in (B). See Results and STAR Methods for more details. Scale bar = 200μm. **(D)** Comparison between the F1 scores and the mean Average Precision averaged over IoU thresholds from 50% to 95% at 5% steps (mAP@50:5:95) between the model described in (A) and a comparable model, Yolov8. **(E)** The *R*^2^ and normalized RMSE (NRMSE, normalized by the mean) for all morphological traits were calculated from the stomatal images in the validation dataset (see Table 1, S2). See Fig. S2 for calculated versus ground truth measurements for all traits.

**Table 1:** List of notation and corresponding descriptions for the 14 stomatal traits calculated in this work.

| Notation | Trait | Description | Unit | Category |
| --- | --- | --- | --- | --- |
| SW | Stomata width | Short axis of the stomata | $\mu\text{m}$ | Size |
| SL | Stomata length | Long axis of the stomata | $\mu\text{m}$ | Size |
| SS | Stomata size | Width by length ($\text{SW} \times \text{SL}$) | $\text{mm}^2$ | Size |
| W/L | Stomata width-to-length ratio | Ratio between width and length, closer to 1 is squarer in shape | unitless | Size |
| SDW | Deviation in stomatal width | Standard deviation of stomata width | $\mu\text{m}$ | Deviation |
| SDL | Deviation in stomatal length | Standard deviation of stomata length | $\mu\text{m}$ | Deviation |
| SDS | Deviation in stomatal size | Standard deviation of stomata size | $\text{mm}^2$ | Deviation |
| SDWL | Deviation in stomatal width-to-length ratio | Standard deviation of stomata size | unitless | Deviation |
| SD | Stomatal density | Number of stomata per image area | Number of stomata per $\text{mm}^2$ | Coverage |
| PSA | Percentage of stomatal area within one image | Covered area of stomata within one image | % | Coverage |
| $g_{smax}$ | Maximum stomatal conductance | The fastest rate at which gases diffuse through fully open stomata (see Eq. 4) | $\text{mol m}^{-2} \text{s}^{-1}$ | Coverage |
| $SE_{ve}$ | Stomatal evenness | The regularity of the distribution of stomata (see Eq. 1), from Liu et al. | unitless, between 0 (clustered) and 1 (even) | Patterning |
| $SD_{iv}$ | Stomatal divergence | The divergence in the distribution of stomata based on their distances from their center of gravity (see Eq. 2), from Liu et al. | unitless, between 0 (close to the center of gravity) and 1 (far from the center of gravity) | Patterning |
| $SA_{gg}$ | Stomatal aggregation | The stomata distribution type (i.e., clustered, random, or uniformly distributed) based on the nearest-neighbor distance (see Eq. 3); from Liu et al. | unitless, between 0 and 2.15 (uniform: $SA_{gg} > 1$; random: $SA_{gg} = 1$; clustered: $SA_{gg} < 1$) | Patterning |

Using a portable microscope with 400× magnification, we obtained a total of 26,893 images of both the abaxial (AB) and adaxial (AD) sides of leaves (see Methods and Table S2). To ensure functional comparability of sampled leaves while accounting for both environment-specific developmental and physiological differences, we selected contrasting leaf ranks that represent the dominant photosynthetic source tissue at each growth stage and experimental setting. Therefore, leaf selection was aligned with the primary phase of carbon assimilation relevant to early vigor or reproductive maturity. In growth chamber experiments, we imaged the third leaf at 3-5 weeks post-transplantation, corresponding to the earliest fully expanded and developmentally stable leaf during early vegetative growth, when plants exhibit high sensitivity to environmental conditions.^43–45^ In the greenhouse experiments, we imaged both the third and sixth leaves, capturing early source activity prior to stress imposition and later-developed source tissue responding to sustained water limitation, respectively (see Methods). In the field experiment, we imaged the flag leaf during the grain-filling phase, due to its significant contribution to yield performance at reproductive maturity.^46,47^

### A high-throughput computational pipeline can accurately identify, classify, and measure stomata from microscopy images

To analyze our images, we constructed a high-throughput computational pipeline to identify and classify objects into one of the following categories: complete stomata, incomplete stomata, blurry complete stomata, blurry incomplete stomata, and hair/trichomes (Fig. 1B). Our proposed framework for stomata detection consists of two semi-supervised learning pipelines: (i) a structural learning pipeline to capture unidirectional alignments of stomata, and (ii) a blurry object learning pipeline to distinguish blurry stomata from trichomes. First, due to developmental patterns, stomata of wheat leaves generally exhibit a pronounced unidirectional alignment pattern, where they tend to appear in rows rather than being randomly distributed. Accordingly, our structural learning pipeline employs an attention mechanism^48^ to model spatial correlations among stomata, leveraging unidirectional alignment as an important structural feature for distinguishing true stomata from false detections. Second, some blurry stomata are difficult to distinguish from plant trichomes, as they present highly similar visual characteristics. Therefore, the blurry object learning pipeline incorporates blurry augmentation to simulate diverse blur patterns observed in real-world images, thereby enhancing the model’s ability to detect blurry objects.

In both pipelines, we employed a teacher-student framework to improve detection accuracy by leveraging unlabeled data, as large-scale manual annotation is impractical and labor-intensive.^49^ The framework operates through the collaborative interaction between a teacher model and a student model. The student model is trained using both labeled stomata and pseudo-labels generated by the teacher model. Meanwhile, the teacher model, updated as an exponential moving average of the student parameters, provides additional supervisory knowledge from unlabeled samples, thereby enabling effective utilization of unlabeled data without requiring additional human annotation (Fig. 1B).

To train our model, we used a manually labeled training dataset of 396 images, which included diverse experimental conditions and genotypes (Table S3). After training, we validated our model using an independent data set that consisted of 240 manually labeled images (see Methods and Table S3) and compared our model against the object detection and classification pipeline Yolov8.^50^ We find that our model performs better at both object detection and classification, particularly with incomplete stomata and hair classification (Fig. 1C–D). Both the F1 and mean average precision (mAP) for our model are close to 1 and 100, respectively, indicating near-perfect object detection and classification. In comparison to the only currently available pipeline trained on high-resolution (×400) wheat images (Rapid Method)^31^, our pipeline can identify the stomata in their images (Fig. S1A), but their pipeline struggles to identify the stomata in our images (Fig. S1B). The Rapid method fails to identify any stomatal detections in 27.9% of our images, and in total, 64.6% of ground-truth stomata were unidentified.

From each image, the coordinates of detected stomata were used to quantify 14 stomatal traits, encompassing both morphological and coverage-related characteristics (see Table 1 and Methods). Eight were calculated using only complete stomata within each image: (1) median stomatal width (SW), (2) median stomatal length (SL), (3) median stomatal width to length ratio (W/L), and (4) median stomatal size (SS), along with their respective standard deviations: (5) standard deviation of length (SDL), (6) standard deviation of width (SDW), (7) standard deviation of width to length ratio (SDWL), and (8) standard deviation of size (SDS). In addition, three coverage-related traits were calculated: (9) stomatal density (SD), i.e., the number of effective stomata per unit area (with “complete” stomata weighted as 1 and “incomplete” stomata as 0.5), (10) percentage of stomatal area (PSA), defined as the ratio of the total weighted stomatal area to the total image area, and (11) maximum stomatal conductance (*g_smax_*, Eq. 1).^11,51^ Finally, three indices were used to characterize the spatial distribution of stomata:^15^ (12) the stomatal evenness index (*SE*_ve_, Eq. 2), (13) the stomatal divergence index (*SD*_iv_, Eq. 3), and (14) the stomatal aggregation index (*SA*_gg_, Eq. 4).

Using the validation dataset, we found that our pipeline accurately calculated ten out of the fourteen selected traits (Fig. 1E, S2). Traits that characterize stomatal size (SW, SL, SS, W/L) show strong correlations with the ground truth dataset (*R*^2^ > 0.7) and very low error (NRMSE < 0.12; Figs. S2A–D). Deviation traits (SDW, SDL, SDS, SDWL) show weak correlations and high error (*R*^2^ < 0.4, NRMSE > 0.2; Figs. S2E–H). Stomatal coverage traits (SD, PSA, *g_smax_*) show very strong correlations and very low error (*R*^2^ > 0.9, NRMSE < 0.1; Figs. S2I–K). Lastly, among the patterning traits, *SD_iv_* also exhibited very strong correlations with low error (*R*² = 0.93, NRMSE = 0.05; Fig. S2L), comparable to the stomatal coverage traits, whereas *SA_gg_* and *SE_ve_* showed strong correlations and low error (*R*² > 0.7, NRMSE < 0.1; Figs. S2M–N). Similar to the detection results, we found that the Rapid method could not accurately quantify most of the stomatal traits from our images, with low *R*^2^ and high NRMSE values for size (Fig. S3A–D), deviation (Fig. S3E–H), coverage (Fig. S3I–K), and patterning (Fig. S3L–N).

After validating our pipeline, we processed a total of 26,645 images and detected 1,452,599 objects, including 858,229 stomatal objects categorized as follows: 270,512 complete, 461,555 blurry complete, 29,662 incomplete, and 96,540 blurry incomplete. Additionally, the pipeline identified 594,370 trichome structures, which we did not analyze for the remainder of this work.

### Ten stomatal traits characterizing stomatal size, patterning, and coverage are highly broad-sense heritable and plastic in response to the environment

We estimated the broad-sense heritability (Eq. 5, *H*^2^) of each trait for each side, environmental condition, and treatment (Figs. 2, S4).^52^ We find that size (SW, SL, SS, W/L), patterning (*SD_iv_*, *SA_gg_*, *SE_ve_*), and coverage-related (SD, PSA, *g_smax_*) traits are highly broad-sense heritable, particularly under field and GC conditions (0.76 < *H*^2^ < 0.95); however, deviation traits (SDW, SDL, SDS, SDWL) are much less heritable for all conditions, suggesting that they are either not genetically controlled or improperly detected (*H*^2^ < 0.3-0.4; Fig. 1E). The high broad-sense heritability observed under GC conditions across treatments suggests that early-stage phenotyping can reliably capture genetic variation that is also expressed at a much later developmental stage under field conditions. Hereafter, we do not consider the less heritable traits (SDW, SDL, SDS, SDWL) and focus our attention strictly on the 10 traits that are the most broad-sense heritable. Most traits under greenhouse conditions have lower broad-sense heritability compared with the field and GC conditions (Figs. 2, S4). Under GC conditions, the abaxial surface generally showed higher broad-sense heritability compared to the adaxial surface. However, this result was reversed under field conditions.

**Figure 2:**
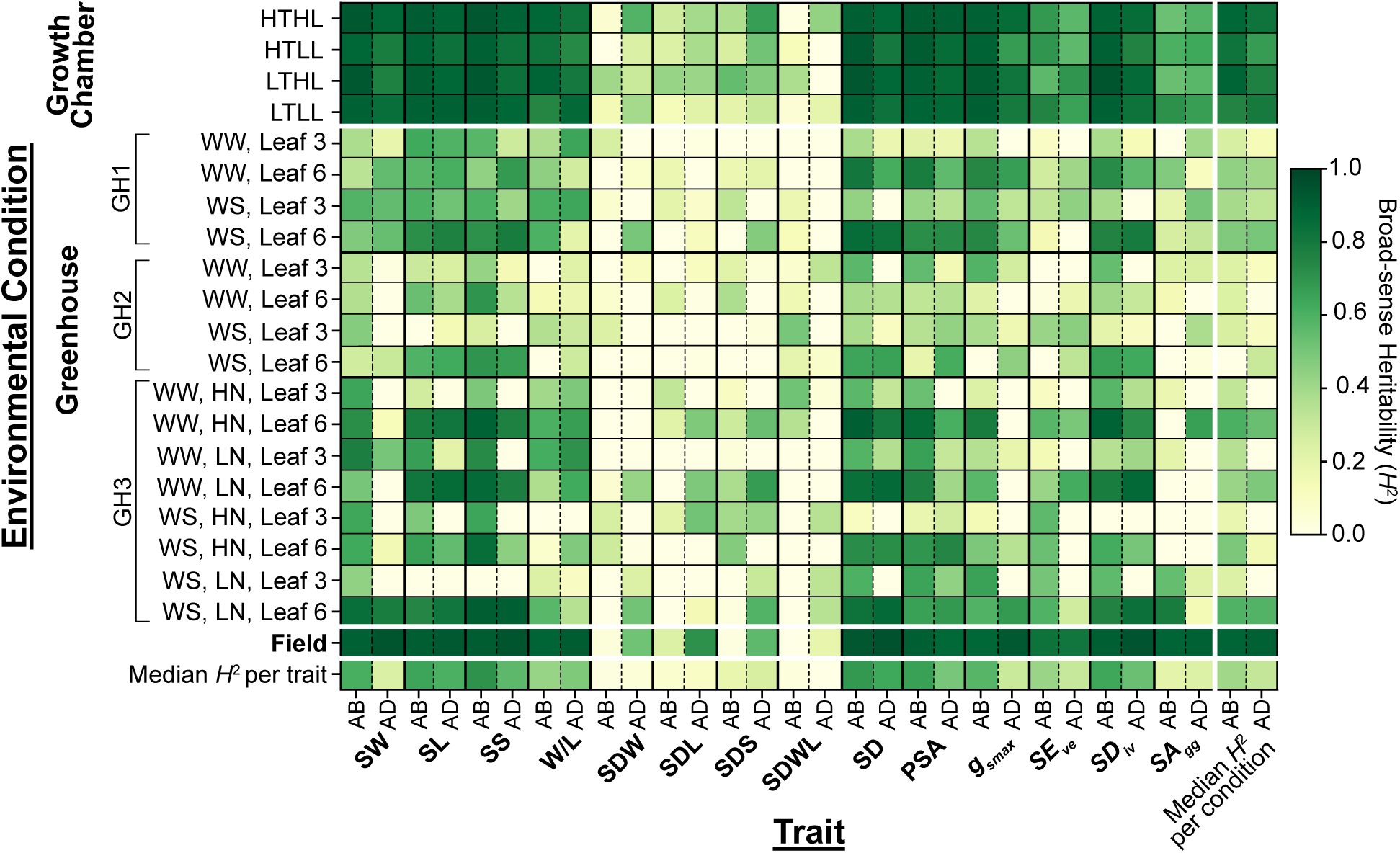
Ten stomatal traits from growth chamber and field experiments are highly broad-sense heritable. Broad-sense heritability (*H*^2^, Eq. 5) for 14 stomatal traits (see Table 2) of abaxial and adaxial leaf sides across various environments: (1) four growth chamber treatments combining high temperature (HT) and low temperature (LT) with high light (HL) and low light (LL); (2) two greenhouse experiments (GH1 and GH2) that assessed the effects of drought stress (WS - water stressed; WW - well-watered) on the third and sixth leaf; (3) one greenhouse experiment (GH3) that assessed the effects of both drought stress (WS - water stressed; WW - well-watered) and nitrogen levels (LN - low nitrogen; HN - high nitrogen levels); and, (4) a field experiment with no additional treatments. The median heritability of each axis is displayed in the final column or row, depending on the calculation. Asterisks denote a broad-sense heritability difference of at least 0.1 between the adaxial and abaxial of a specific trait and condition, with the asterisk placed on the higher value’s spot.

To assess the various effects on stomatal traits, we compared the value of our calculated traits across growth conditions, resource availability, and leaf side (Figs. 3, S5). We found that the stomatal density (SD) of the adaxial side was consistently higher than that of the abaxial side across all tested conditions, reaching its highest value under field conditions and lowest under low temperature low light (LTLL) environment (Fig. 3A). Next, stomatal size traits are maximized under low light conditions; stomatal width (SW) and size (SS) are maximized under low temperature but high light conditions (LTHL) environment, while stomatal length (SL) was maximized under either low temperature and low light (LTLL, adaxial) or low temperature high light (LTHL, abaxial) environments (Figs. 3B–C, S5A). Stomatal width (SW) showed more significant differences between adaxial and abaxial sides than stomatal length (SL), implying that stomatal width is more plastic between leaf sides than length (Fig. 3B–C). We also found that stomata on flag leaves from field experiments tend to be squarer in shape (W/L ratio closer to 1) than the third leaves grown in high light conditions (Fig. S5B). Other coverage metrics, such as PSA and *g_smax_*, are significantly higher on the adaxial side across all conditions and are maximized under field and HTHL conditions (Figs. S5C–D). Lastly, patterning traits have different responses to the environment (Figs. S5E–G). Stomatal evenness (*SE_ve_*) is relatively homogeneous across environments, whereas divergence (*SD_iv_*) and aggregation (*SA_gg_*) are significantly higher on the abaxial side for most environments, and, in particular, in GC and field conditions.

**Figure 3:**
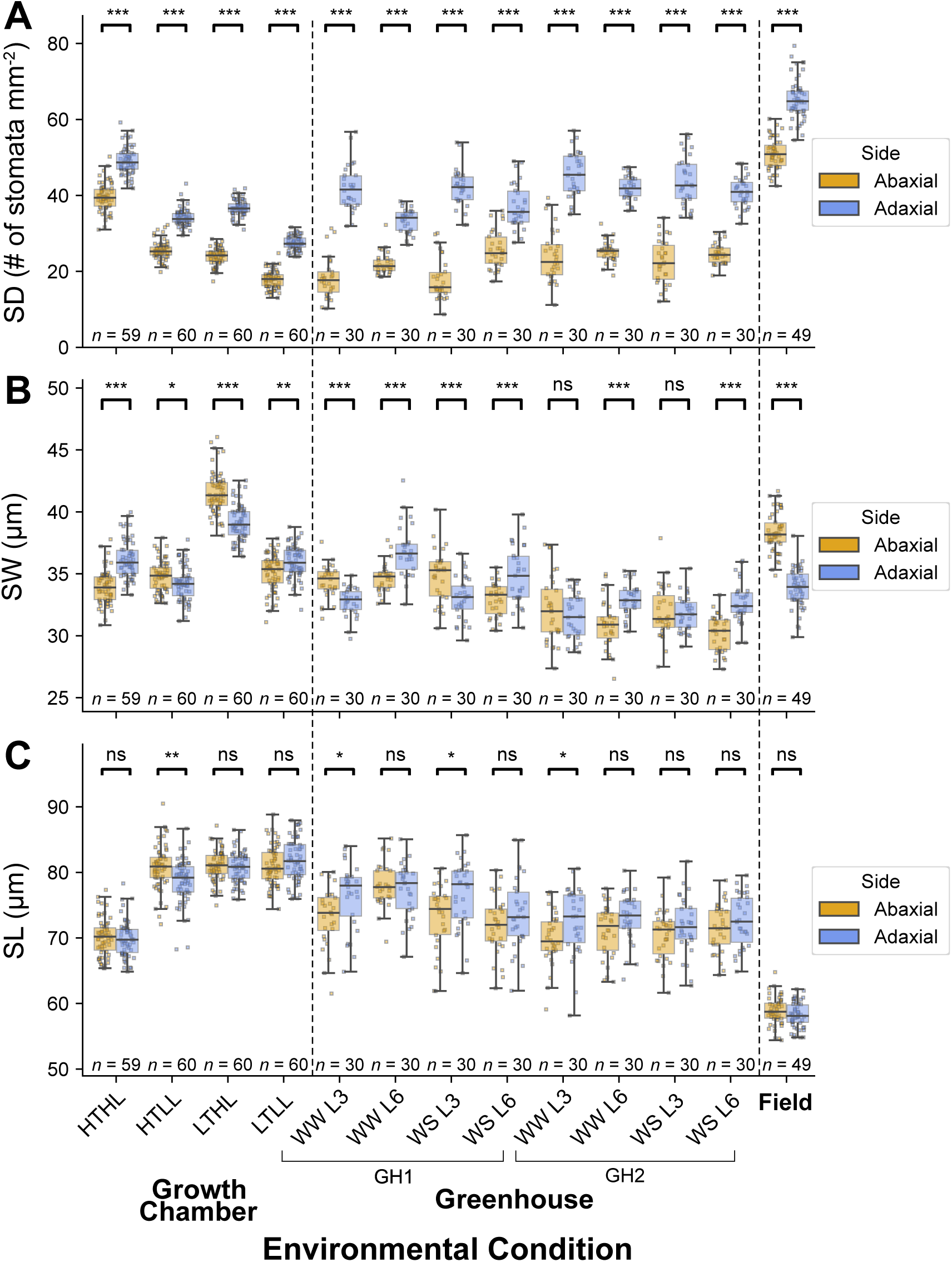
Stomatal traits show environment- and leaf-side-dependent differences in genotypic and plastic responses. Comparison between **(A)** stomatal density (SD, number of stomata per mm^2^), **(B)** stomatal width (SW, μm), and **(C)** stomatal length (SL, μm) of the abaxial and adaxial sides of leaves in winter wheat cultivars grown under various environmental conditions: (1) four growth chamber treatments combining high temperature (HT) and low temperature (LT) with high light (HL) and low light (LL); (2) two greenhouse experiments (GH1 and GH2) that assessed the effects of drought stress (WS - water stressed; WW - well-watered) on the third leaf (L3) and sixth leaf (L6); and (3) a field experiment with no additional treatments. Each point represents the mean trait value for a single wheat cultivar in a specific environmental condition. A pairwise Student’s *t*-test was conducted for each adaxial/abaxial pair with the following notation: * - *p* < 0.05, ** - *p* < 0.01, and *** - *p* < 0.001. See Figs. S3 for additional traits.

We then explored potential coordination in stomatal development between the two leaf surfaces by comparing morphological and patterning traits on the abaxial and adaxial sides of field and growth chamber (GC) conditions (Fig. 4). We find that there is moderate to strong coordination between leaf sides across all environments in traits related to size (SS, Fig. 4A), coverage (SD, Fig. 4B), and patterning (*SD_iv_*, Fig. 4C). We also observe some traits that have moderate coordination between leaf sides in GC conditions but not in the field (SL and PSA, Figs. 4D–E) and vice-versa (*SA_gg_* and *SE_ve_*, Figs. 4F–G). Other traits have more nuanced interpretations, such as having stronger coordination in HTHL environments (*g_smax_*, Fig. 4H) or weaker coordination in LTHL environments (SW and W/L, Figs. 4I–J). These contrasting patterns highlight the environment-dependent regulation of stomatal development between leaf sides and the underlying genetic diversity shaping these responses to environmental conditions.

**Figure 4:**
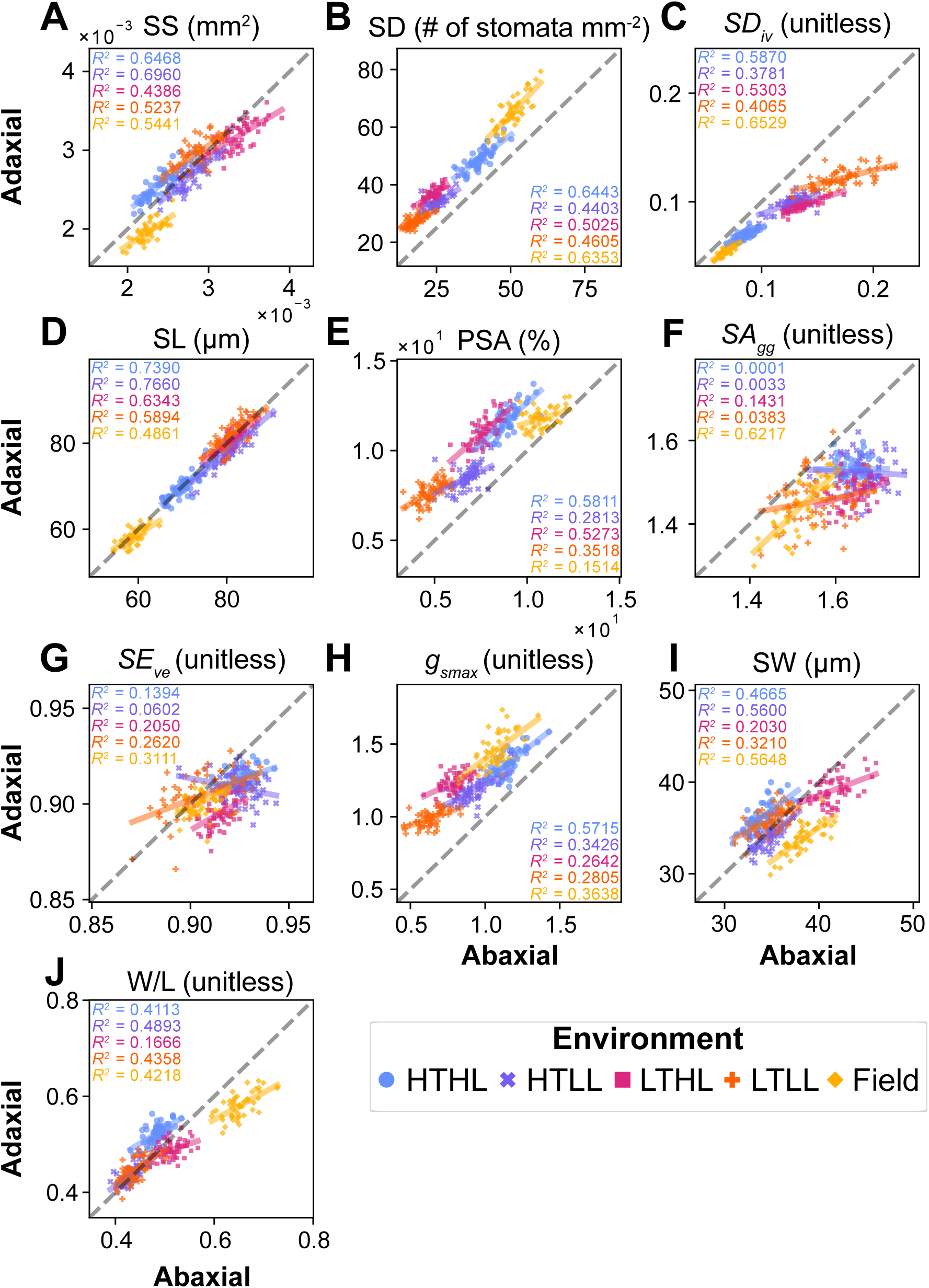
Analysis reveals environment-dependent coordination of stomatal traits between abaxial and adaxial sides of the leaves. Various stomatal morphological and patterning traits are calculated for both the abaxial (horizontal axis) and adaxial (vertical axis) sides of the leaves in various environmental conditions. The traits are **(A)** stomatal density (SD), **(B)** stomatal width (SW), **(C)** stomatal length (SL), **(D)** stomatal size (SS), **(E)** stomatal width-to-length ratio (W/L), **(F)** percentage of stomata area (PSA), **(G)** stomatal conductance (*g_smax_*), **(H)** stomatal evenness (*SE_ve_*), **(I)** stomatal divergence (*SD_iv_*), and **(J)** stomatal aggregation (*SA_gg_*), See Table 2 for information about each trait. Results are shown across field conditions (yellow diamonds) and the four growth chamber experiments: high temperature with high light (HTHL, blue circles), high temperature with low light (HTLL, purple crosses), low temperature with high light (LTHL, red squares), and low temperature with low light (LTLL, orange pluses). Each point represents one cultivar, and the dashed line represents no difference between adaxial and abaxial. Regression lines are plotted for each environmental condition, and *R*^2^ values are colored and provided in each subplot (in order from top to bottom: HTHL, HTLL, LTHL, LTLL, and field).

### Stomatal traits show significant effects of genotype and leaf side, particularly under field and growth chamber conditions

We wanted to quantify the impact that environmental conditions and plant traits had on each of the broad-sense heritable stomatal traits. To this end, we used a linear mixed model to quantify the effect of several important factors tested in our work and their interactions (see Methods): genotype (G), leaf side (LS), temperature (T), light (L), leaf rank (LR), water treatment (W), and nitrogen treatment (N). Almost all traits we analyzed show significant effects of genotypes and leaf sides (i.e., abaxial/adaxial) under field and GC environments (Fig. 5A-B). The effects of temperature, light, and water stress on SW and SL were significant across most conditions, except for SL in GH2 and the effect of water stress on SW and SL in GH3 (Fig. 5). Although the traits studied under greenhouse environments were less broad-sense heritable (Fig. 2), most of the traits showed significant genotypic differences except evenness and aggregation (Fig. 5C–E). Despite the limited numbers of cultivars, nitrogen (N) significantly affected SL and SS in greenhouse conditions, highlighting the high plasticity of these morphological traits in response to nutritional variation (Fig. 5E). The interactions between genotype and leaf side (G:LS) were also significant, except in GH2 (Fig. 5D). Despite the significant influence of light and leaf sides on SW, SL and W/L, their interaction was not significant, particularly in GC conditions, suggesting that irradiance-driven stomatal plasticity operates at the whole-leaf level ensuring stable diffusion efficiency and transpiration control (Fig. 5B).

**Figure 5:**
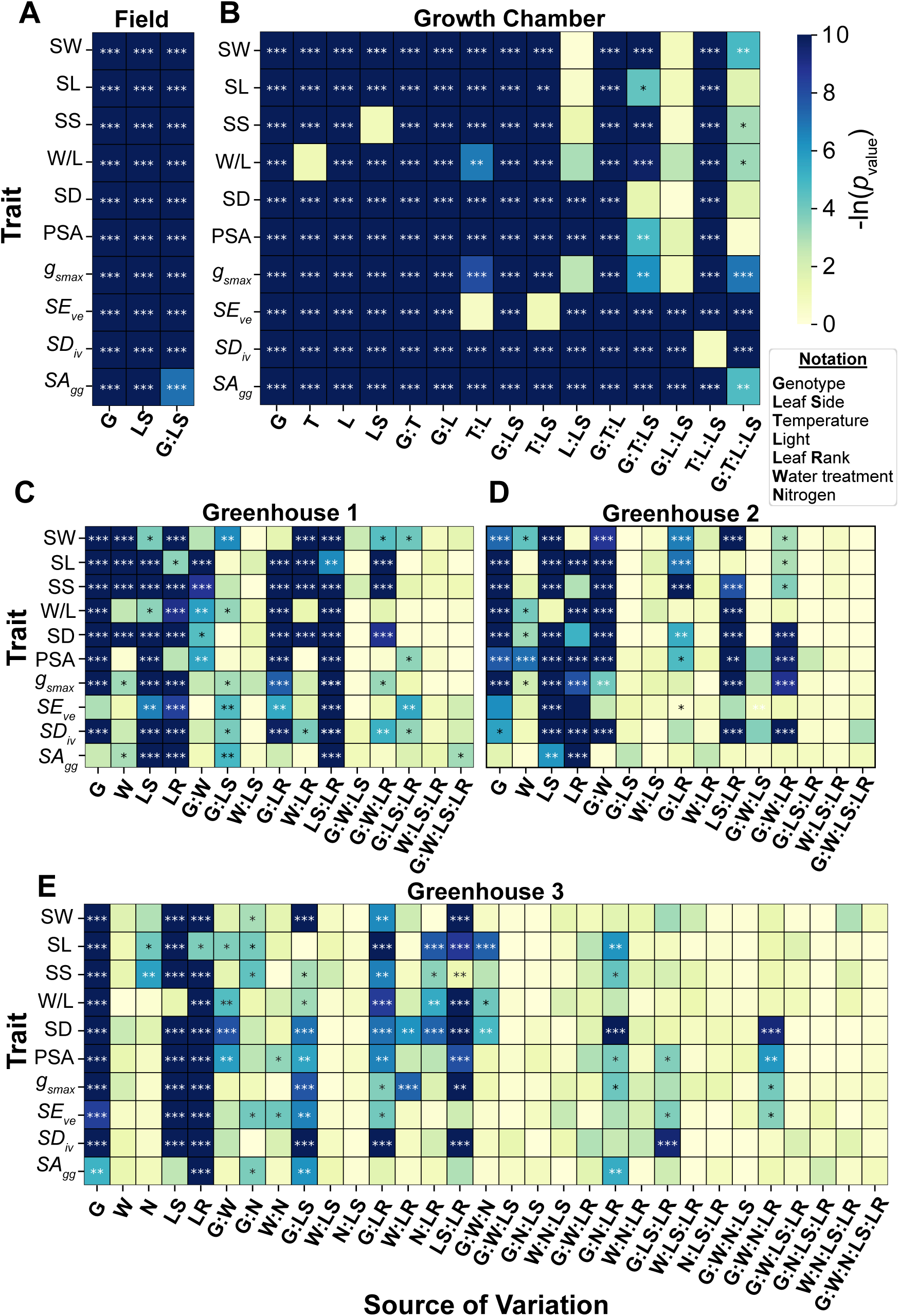
Stomatal traits show significant effects of genotype and leaf side, particularly under field and growth chamber conditions. Heatmaps showing statistical significance of the effect of the following factors on the 10 heritable stomatal traits (see Table 2, Fig. 2, and STAR Methods): genotype (G), leaf side (LS), light (L), temperature (T), leaf rank (LR), water treatment (W), and nitrogen (N). Results are shown across different environments: **(A)** field, **(B)** growth chamber, and greenhouse experiments **(C)** GH1, **(D)** GH2, and **(E)** GH3 (see Table 1). For each condition and trait, a one-way ANOVA model was fit, and the natural logarithm of the corresponding *p*-value is plotted. Significant results are represented by asterisks: * - *p* < 0.05, ** - *p* < 0.01, and *** - *p* < 0.001.

Interestingly, under field conditions with higher light intensity, stomata on flag leaves tend to be squarer in shape, with a W/L ratio close to 1 (0.6 for adaxial and 0.7 for abaxial side, respectively, Fig S5B). This was followed by other high light conditions—HTHL (0.52 and 0.48 for AD and AB, respectively) and LTHL (0.5 and 0.51 for AD and AB, respectively, Fig. S5B). While the temperature effect on W/L was not significant, light and its interaction with temperature influenced W/L and stomatal geometry, indicating that the plasticity of stomatal shape (as reflected by the W/L ratio) in response to varying light and temperature conditions is driven by interactive rather than additive effects (Fig. 4B). Surprisingly, both elevated temperature and high light significantly reduced stomatal divergence on both leaf surfaces, with the lowest values observed under field conditions and in the HTHL treatment (Fig. S5F), contrasting with their effect on evenness (Fig. S5E).

Interactions between environmental factors and all stomatal traits were generally insignificant under greenhouse conditions (Fig. 4C-E), particularly for water and nitrogen availability, most likely because greenhouse traits were less broad-sense heritable (Fig. 2). In contrast, climatic variables such as light and temperature elicited more consistent and structured interactions (Fig. 4C–E). These findings coincide with our findings on water use efficiency (WUE), as we found that – in general – stomatal size and density do not influence WUE when nitrogen or water stresses are applied (Fig. S6).

### Growth chamber experiments on the third leaf capture the genetic variation of stomatal conduction on flag leaves in the field

It has been frequently reported that the results in the lab conditions can not represent the results from the field.^53,54^ To this end, we investigated the comparability of stomatal traits between the field and controlled environments. In growth chamber (GC) and greenhouse (GH) conditions, stomatal morphology was phenotyped on the leaves in the early developmental stages, typical of the phenotyping procedure in the lab environments. Correlation analysis was performed on ten broad-sense heritable traits. GH3 was excluded from the analysis due to the small sample size (*n* = 10).

We found that the correlation of stomatal traits between controlled (GC) and field conditions were stronger than between GH conditions, highlighting the role of genetic variance in ensuring trait repeatability across environments (Fig. 5). Size (SW, SL, SS) and coverage (W/L, SD, PSA, *g_smax_*) traits in GC conditions correlated well with the field, highlighting the potential of using GC conditions to phenotype stomatal morphology. There were significant differences in how leaf side played a role in these correlations, as the HTHL condition had a similar median correlation value between leaf sides (abaxial: 0.454, adaxial: 0.461), whereas HTLL had a statistically significant difference between leaf sides (abaxial: 0.477, adaxial: 0.248; Fig. S7). Some correlations are leaf side-dependent, as across almost all environmental conditions, *g_smax_* and PSA correlated more strongly with the field on the abaxial rather than the adaxial side, particularly in GC conditions (Fig. 5).

In contrast, correlations between field and GH conditions were generally weaker and less significant than GC conditions (Fig. 5). As well, broad-sense heritable traits expressed across environments did not always correlate strongly, highlighting an environment-dependent phenotypic plasticity. For instance, in the GH1 water stressed (WS) condition, stomatal length (SL) and stomatal size (SS) exhibited high broad-sense heritability in the 6th leaf (Fig. 2). However, they were weakly correlated with field measurements (abaxial - SL: *r* = 0.03, SS: *r* = 0.03; adaxial - SL: *r* = 0.06, SS: *r* = 0.27; Fig. 5). From the patterning traits, we found that only *SA_gg_* positively correlated with the field, whereas both *SE_ve_* and *SD_iv_* either were negatively correlated or did not correlate at all. These findings demonstrate that water availability modulates genotype-specific plasticity of stomatal traits, thereby limiting the predictive value of measurements under greenhouse conditions for field performance.

### Breeding maintains the diversity of stomatal traits and increases the transpirational potential under field and high temperature conditions

To assess whether breeding history has influenced stomatal traits, we estimated the relative breeding progress (RBP) for the ten broad-sense heritable traits measured under both field and growth chamber conditions (see Methods).^55^ An RBP value of 1 indicates no change over time. Strikingly, we found that under all growth chamber and field conditions, breeding has increased the percentage of stomatal area (PSA) on both sides of the leaf. Under high temperature and field conditions, stomatal size (SS) and stomatal width (SW) have increased, slightly more pronounced on the adaxial side. Also, maximal stomatal conductance (*g_smax_*) increased only under growth chamber conditions. Patterning traits such as stomatal divergence (*SD_iv_*) decreased with breeding, most likely due to the increase in PSA during the same period. Most other traits showed minimal to no evidence of selection in these environmental conditions (Fig. 7A). As well, breeding progress was more pronounced on the adaxial surface than on the abaxial surface (Fig. 7A). These findings suggest that modern cultivars possess plasticity to enhance leaf cooling capacity or photosynthetic rate under high temperatures (25-30°C). Furthermore, by using reverse genomic selection,^56^ we detected reverse genomic selection signals that were generally consistent with the observed relative breeding progress, showing that the observed phenotypic trends across breeding history are also supported at the genomic level by corresponding changes in allele frequencies (Fig. 7B; see Methods).

### German and non-German wheat varieties generally show similar genetic diversity in stomatal traits

In our panel of sixty winter wheat cultivars, fifty of the studied cultivars represented the breeding history of winter wheat in Germany between 1966 and 2018 (the breeding panel), and ten represented exotic cultivars (the exotic panel). We aimed to see if there were differences in stomatal adaptation in growth chamber conditions between these two panels (Fig. S8). We find that, in general, there are few differences between the stomatal traits of the breeding and exotic panels across all traits and environmental conditions. However, there are slight differences between the two groups, mainly under LTLL conditions. We found an increased percentage of stomatal area (PSA) in the exotic panel on both sides of the leaf, mainly attributed to the increase in stomatal width (SW) and, as a byproduct, the width-to-length ratio (W/L), combined with an insignificant difference between stomatal density (SD, Fig S8). Therefore, our data suggests that there is little significant difference between the breeding and exotic panels, apart from a few, but significant differences in some traits under low light and temperature conditions.

## Discussion

In this study, we analyzed stomatal morphological traits in 60 winter wheat cultivars (Table S1) using a multi-environment dataset of 26,893 image observations from field, growth chamber, and greenhouse experiments generated by a customized automatic detection pipeline. This dataset allowed us to assess the plasticity of stomatal morphology and patterning in response to light, temperature, drought, and nitrogen (Table S2). Four main findings emerged: (i) stomatal traits were largely plastic in response to environmental conditions and showed genotype-specific responses (Fig. 3, S5); (ii) traits on the 3rd leaf under growth chamber conditions reliably captured genetic variance expressed at a much later developmental stage on the flag leaf in the field (Fig. 6, S7); (iii) the upper leaf surface had consistently higher stomatal density, area, and maximum conductance, while systematic proper patterning and spacing were more found on the lower surface (Fig. 4); and (iv) breeding selection favored stomatal area (PSA) over density (SD) across field and controlled environments (Fig. 7).

**Figure 6:**
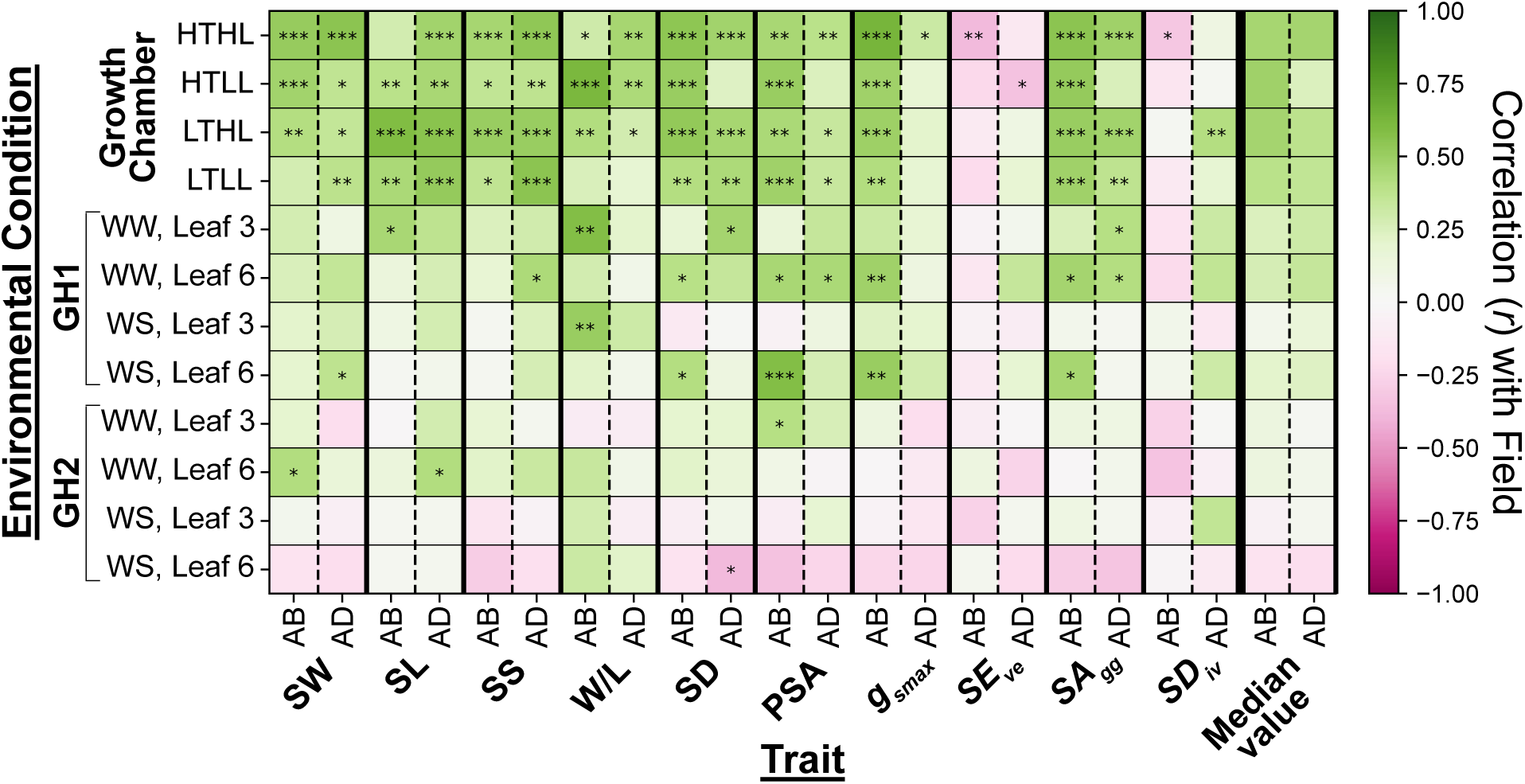
The stomatal traits of third leaves grown in growth chambers correlate strongly with flag leaves grown in the field. A heatmap of the Pearson correlation coefficients (*r*) of 10 heritable stomatal traits (see Table 1) between field and controlled environments (greenhouse and growth chambers), where pink represents a negative correlation and green represents a positive correlation with the field. The controlled environments include 1) four growth chamber treatments combining high temperature (HT) and low temperature (LT) with high light (HL) and low light (LL), and 2) two greenhouse experiments (GH1 and GH2) that assessed the effects of drought stress (WS - water-stressed; WW - well-watered) on the third and sixth leaves. Corresponding boxplots for each environmental condition are shown in Fig. S4. Significant results are represented by asterisks: * - *p* < 0.05, ** - *p* < 0.01, and *** - *p* < 0.001.

**Figure 7:**
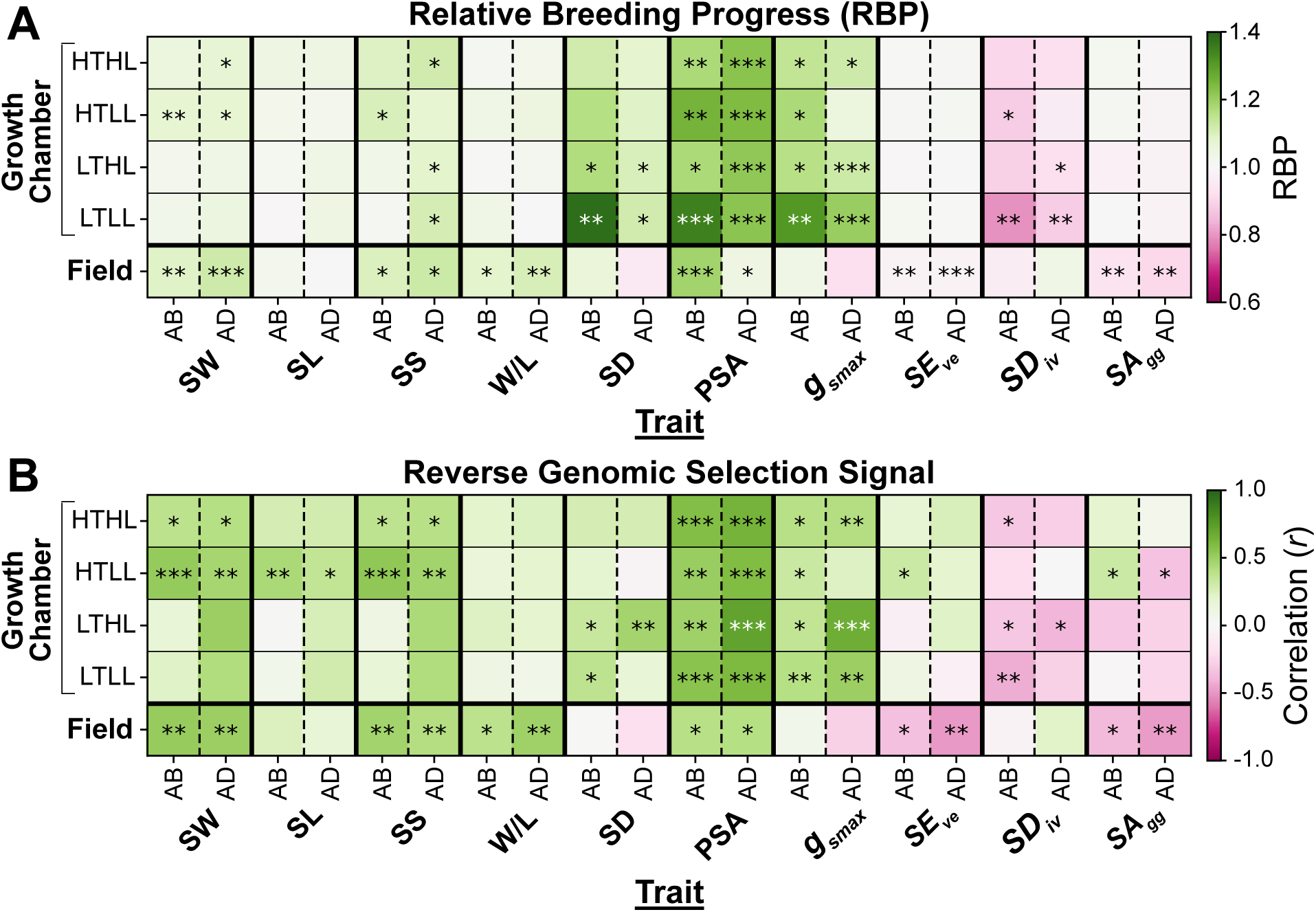
Breeding selection favored stomatal area (PSA) over density (SD) across field and controlled environments. **(A)** Relative breeding progress (RBP) and **(B)** reverse genomic selection signal of 10 heritable traits. Results are shown across field conditions and the four growth chamber experiments: high temperature with high light (HTHL), high temperature with low light (HTLL), low temperature with high light (LTHL), and low temperature with low light (LTLL). Stomatal traits include stomatal width (SW), length (SL), size (SS), width-to-length ratio (W/L), stomatal density (SD), percentage of stomatal area (PSA), maximum stomatal conductance (*g_smax_*), evenness (*SE_ve_*), divergence (*SD_iv_*), and aggregation (*SA_gg_*). Statistically significant correlations are indicated within the colored squares: *** - *p* < 0.001, ** - *p* < 0.01, * - *p* < 0.05; empty squares denote non-significant correlations.

### A promising pipeline for quick quantification of stomatal traits in lab and field environments

Efficient and accurate quantification of stomatal traits is critical for advancing research on plant gas exchange and morpho-physiological responses while also bridging the gap between genomics and phenomics.^7,13,21,57–59^ Recent efforts have focused on integrating user-friendly high-resolution lenses (e.g., 400×) with computer vision pipelines to enable rapid and automated stomatal phenotyping.^28,30,31,40,60^ However, some of these pipelines struggle with processing and identifying stomata on leaves that are not completely flat (Figs. S1, S3). Our pipeline addresses this limitation by effectively distinguishing between complete, blurry, and incomplete stomata, improving the accuracy of stomatal size quantification, regardless of the leaf morphology (Figs. 1, S1, S2, S3). Because of this flexibility in the pipeline, an experienced user can acquire each image within approximately 15 seconds, greatly facilitating throughput. While our current model can be easily used to detect stomata in similar species (e.g., barley), training the model for new, more distantly related species typically does not require many images and can be done relatively quickly with an annotated dataset of 50 to 100 images of sufficient variety.^61^ However, the pipeline may require significant retraining when applied to images acquired at different resolutions or magnification levels (e.g., 50× or 200×), as variations in image scale can affect model performance. Sample preparation is another important consideration. Leaves with highly waxy surfaces, particularly the abaxial side of flag leaves, may require gentle cleaning before imaging to minimize image distortion, improve stomatal visibility, and ensure accurate detection, which can be prohibitively time-consuming.

Using our pipeline, we uncovered substantial genetic variation in stomatal traits and their plasticity across environments (Figs. 3, 5, S5). Such plasticity can lead to broad-sense heritable trait expression within individual environments (Fig. 2), yet weak correlations across environments (Fig. S7), echoing the longstanding challenge of translating growth chamber findings to field performance.^54^ Notably, stomatal traits measured on third leaves under growth chamber conditions correlated more strongly with field-grown flag leaves than typically reported (Fig. 6).^54,62^ This suggests that stomatal development is under tight genetic control throughout the life cycle, whereas trait variation is largely environment-driven.^63^ This may also explain how both indoor phenotyping of adaptive traits, such as canopy stomatal conductance, can enable genomic prediction of field performance,^41,64^ and the need to reassess experimental strategies to improve screening traits under controlled environments should be considered.^65^ Our results support the potential use of seedling phenotyping under stressful, field-like conditions to dissect the genetic basis of stomatal traits underpinning stress tolerance in the field, where multiple physiological processes and developmental stages interact.^66^

### Stomatal size traits exhibit strong responses to selection and environmental variation

Domestication and breeding have generally reduced abaxial stomatal density with the increase of stomata size in cultivated species compared with their wild relatives.^67^ Although stomatal traits have not been directly targeted by breeders, yield improvement in German winter wheat has been associated with a higher proportion of whole stomatal area on both sides of the leaf, primarily through increased stomatal size, while density remained unchanged (Fig. 7A). These phenotypic trends were largely supported by the reverse genomic selection signals, which showed consistent positive selection signals across environments for PSA, as well as predominantly positive signals for stomatal width and size (Fig. 7B). The consistency between the phenotypic relative breeding process and the reverse genomic selection signal supports the view that increases in stomatal area have been a consequence of selection during modern wheat breeding, even though stomatal traits were not direct breeding targets. In contrast, rice breeding for yield improvement has favored reduced density on flag leaves.^68^ These contrasting trajectories suggest that different grass species, or their respective cultivation practices, employ distinct strategies to optimize photosynthetic capacity and water-use efficiency in support of yield gains under stress conditions.

Unlike stomatal width, size, density, and PSA, stomatal length remained notably stable during breeding (Fig. 7A). This pattern was also reflected by the comparatively weak reverse genomic selection signals detected for this trait (Fig. 7B). This stability highlights a strong developmental constraint, suggesting that stomatal length is a highly conserved anatomical trait that is largely unresponsive to indirect selection for yield improvement. In addition to the potential relevance of PSA, plasticity of size-related traits was more pronounced and differed significantly across size-related traits (SW, W/L, and PSA) between the genetic panels (German and non-German cultivars), particularly under low temperature and low light conditions, while density remained unchanged between the genetic panels (Fig. S8). These findings suggest that within the same condition, the variation in stomatal plasticity among genetic pools is expressed primarily through morphological adjustments of stomatal size rather than changes in stomatal density, unlike *Arabidopsis thaliana*, where variation of density was higher than size among large genotypic variation.^69^

The coordinated functional responses between the adaxial and abaxial leaf surfaces are not governed by light alone; additional factors, such as intercellular CO_2_ and mesophyll-driven signals, play critical roles.^70–73^ One of these factors is the anatomical characteristics of stomata, such as stomatal density, size (guard cell length), and patterning.^4–10^ The finding of a nearly identical 1:1 ratio for stomatal length between surfaces on the 3rd leaf under contrasting temperature and light levels, as well as on the flag leaf under field conditions, suggests that this anatomical trait serves as a coordinated structural anchor across varying environmental conditions (Fig 4D). Such symmetry in stomatal length may contribute to the similar stomatal conductance (*g_smax_*) kinetics observed during step changes in PPFD and could be a stronger determinant of coordinated stomatal behavior than the more dynamic variation in pore width.^10^ Our results are consistent with previous studies showing that higher stomatal density, stomatal area, and maximum stomatal conductance on the adaxial surface of wheat leaves may support greater photosynthetic capacity and enhanced evaporative cooling, thereby helping to maintain favorable leaf temperatures for photosynthesis. In contrast, the lower leaf surface exhibited more systematic stomatal patterning and spacing, which may facilitate efficient gas exchange and CO₂ diffusion as previously proposed.^10,16,74–76^

### Targeting stomatal traits and their plasticity to design climate-resilient crops

Field trials that are complemented with enhanced phenotyping methods under controlled conditions produce reliable and high-quality data for breeders^41,64,77^ because of the interactions between genotype and environment, as shown in stomatal conductance and morphology^63,78^ along with our study (Fig. 5). Our comprehensive analyses provide interesting observations that could be used to design climate-resilient cultivars. For example, interactions of nitrogen and water stress with leaf side were not significant across all stomatal traits (Fig. 5E), in contrast to the significant interactions of temperature and light with leaf side (Fig. 5B). This indicates that the spatial modulation of stomatal traits across leaf surfaces is driven primarily by climatic variables (e.g., light, temperature) rather than by soil-related factors (e.g., water and nitrogen availability). We also observed significant interactions between genotype and leaf side (Fig. 5A–B), indicating both functional asymmetry between adaxial and abaxial surfaces and genotypic plasticity in the deployment of leaf surface traits in response to the environment.^79,80^

Our finding that stomatal traits can be highly broad-sense heritable within environments (Fig. 2) but more weakly correlated across environments (Fig. 6) highlights genotypic differences in stomatal developmental plasticity. This may partly explain why such traits have rarely been direct breeding targets, as cultivar evaluation typically occurs across diverse environments.^62,81^ The lack of consistent selection on stomatal traits may also reflect recent evidence that modern cultivars exploit multiple physiological pathways to achieve high and stable yield,^82^ implying diverse water-use strategies. For example, the cultivar Patras, characterized by fewer tillers and early flowering, combines low stomatal density with large stomatal size and shows no improvement in water-use efficiency under drought, as do most of the cultivars explored in this study (Fig. S6). Nevertheless, this trait combination might stabilize yield through wide stomatal apertures that support photosynthesis and enhanced pre-anthesis carbohydrate storage in stems, especially when water is not the prevailing limiting factor in the field. Thus, high-throughput approaches for quantifying stomatal morphology could provide breeders with a more knowledge-driven basis for selecting cultivars with environment-specific strategies for yield stability.

## Resource availability

### Lead contact

Requests for further information and resources should be directed to the lead contact, Tsu-Wei Chen.

### Materials availability

This study did not generate new, unique reagents. Requests for all materials used in this study should be directed to the lead contact, Tsu-Wei Chen.

### Data and code availability

All data presented in the study are publicly available in a Zenodo data repository (10.5281/zenodo.20609511).

The code for all the programs in this paper, including the stomatal identification and trait quantification, can be found in our GitLab repository, https://scm.cms.hu-berlin.de/intensive-plant-food-systems-public/2026-mabrouk-stomatal-phenotyping.

Any additional information required to reanalyze the data reported in this work is available from the lead contact upon request.

## Acknowledgements

We thank Deutsche Forschungsgemeinschaft (German Research Foundation, DFG) for funding this study. T.-W.C. was funded under “Emmy Noether Programm”, project number 442020478. T.-W.C., R.J.S., A.S., and E.H. were funded by DFG under project numbers 518863370, 518783157, 518913298, and 518914346, respectively. We also thank Constantin Schmidt, Marc Duttlinger, Isabel Dangus, Wen-Ting Chen, Han-Wen Hsu, Marie-Christin Grundmann, and Burak Arinalp for their efforts in manual annotation of stomata and image processing.

## Author contributions

T.-W.C. conceived the study; M.M. and E.V.A. collected the data; J.-A.L. and Y.H. developed the automatic stomatal detection pipeline under the supervision of F.-J.W.; M.M., N.J.R., and T.-C. W. analyzed the data and drafted the manuscript with inputs from T.-W.C., L.F., B.W., and R.S. conducted and designed the field experiment; E.G., G.W., and A.S. conducted and designed the greenhouse experiments. E.H. and A.M. conceived the genetic analyses. M.M. and T.-W.C. selected the genotypes; all authors contributed to the editing of the paper.

## Methods

### Plant materials

Sixty winter wheat cultivars (*Triticum aestivum* L.) were used to demonstrate the genotypic variation in responses to light and temperature. Fifty of the studied cultivars represent the breeding history of winter wheat in Germany (breeding panel), while ten represent exotic cultivars (exotic panel). For the breeding panel, the selection showed the representativeness of breeding history in terms of i) the range of breeding history, ii) breeding progress in grain yield, and iii) genetic diversity (Table S1).^42^ For the exotic panel, we selected the 10 most genetically diverse cultivars from a collection of 24 cultivars using genome-wide SNP profiles and clustering.^55,83^ In clusters with more than one cultivar, the cultivar with the highest plasticity in tiller number, specific leaf area, plant height, and shoot dry weight, in response to planting density, was chosen.^84^

### Plant growth conditions: growth chambers

Growth chamber experiments (Weisstechnik, fitotron HGC1514, Lindenstruth, Germany) were conducted at both high and low light intensities (458 ± 41 and 49 ± 3 μmol m^−2^ s^−1^ photosynthetic photon flux density, referred to as HL and LL, respectively) with high and low temperature regimes (31.3 ± 1.5/28.0 ± 1.2°C and 15.9 ± 1.2/12.6 ± 1.4°C for day/night temperatures, referred to as HT and LT, respectively). After the development of the third leaf, the most homogeneous seedlings were selected and randomized into pots (3.5 × 3 × 16 cm) filled with bedding substrate (Substrat 1, Klasmann-Deilmann GmbH, Geeste, Germany) at a planting density of 180 plants per m^2^. The transplants were grown under a 12-hour light/dark cycle. Temperature was maintained with minimal spatial heterogeneity using sensors (TV-4510 Tinytag View 2, BMC SOLUTIONS, Germany), distributed at canopy height level along the left, middle, and right sections of the GC. Day/night water vapor pressure deficit was maintained at 2.14/1.7 and 0.82/0.63 kPa at HT and LT, respectively.

The experiment that was used for the validation dataset for the stomatal pipeline was conducted in growth chambers in which plants were exposed to a high light and controlled temperature regime of 22.2 ± 1.0°C/12.6 ± 1.2°C for day/night, respectively. The planting and transplantation were the same as the other growth chamber experiments. The day-night water vapor pressure deficit was maintained at 1.19/0.93 kPa, respectively.

### Plant growth conditions: greenhouses

Greenhouse (GH) experiments were conducted using the multi-sensor, high-throughput gravimetric platform PlantArray® (Plant-DiTech LTD, Julius Kühn Institute, Quedlinburg, Germany) to assess the effects of drought and nitrogen availability on water use efficiency and stomatal morphology of the breeding panel. Due to space limitations on the PA platform, the 50 genotypes were tested in two sequential trials. The first trial (GH1) included 30 genotypes, while the second trial (GH2) tested the remaining 20, along with 10 overlapping genotypes from the first trial. These 10 genotypes were selected based on their water-use efficiency (WUE) performance from GH1. WUE is defined as the plant biomass produced relative to the amount of water used. Out of the biomass dry weight gained at the end of the experiment by harvesting the above-ground leaf material and the cumulative transpiration of the plants during the whole growth period, the water use efficiency is calculated (g kg^−1^ H2O). The cumulative transpiration is defined as the sum of the water lost by plant transpiration in all the days pre-harvest. WUE was compared between twelve cultivars with contrasting stomatal density and size under both control and drought treatments during the greenhouse experiments (GH1, GH2, and GH3; Table 1). The selected twelve cultivars were classified into five groups: high stomatal density with large stomatal size (cultivars Forum and RGT Reform), low stomatal density with large stomatal size (cultivars Intro and Patras), intermediate stomatal density with medium stomatal size (cultivars Meister, KWS Santiago and Nordkap), medium stomatal density with low stomatal size (cultivars Caribo, Benno, and Saturn), and medium stomatal density with large stomatal size (cultivars Biscay and Paroli). In the third trial (GH3), four high WUE, two average WUE, and four low WUE cultivars were selected and tested again with an additional nitrogen treatment (Table S1).

Two levels of nitrogen treatment were considered in GH3: low nitrogen (N-low) and high nitrogen (N-high). At sowing, all pots contained 1.62 g of nitrogen derived from the growing medium CL ED73. The N-high treatment corresponded to 150% of the soil nitrogen level and therefore received an additional 0.81 gN, resulting in a total nitrogen supply of 2.45 g per pot. This additional nitrogen in the N-high treatment was applied as ammonium nitrate (NH_4_NO_3_; Carl ROTH GmbH + Co. KG, Karlsruhe, Germany) in two equal applications of 0.405 g per pot. The first ammonium nitrate application was conducted before inducing drought stress, and the second was applied one week later. Each application was dissolved in 50 mL of distilled water. The N-low treatment received no additional nitrogen beyond the soil background (1.62 g N per pot). Pots assigned to the N-low treatment received an equivalent volume of distilled water at each application time. The amount of ammonium nitrate applied was calculated based on its nitrogen content (35.0% of its molar mass, 80.04 g mol⁻¹). Accordingly, 1.157 g NH_4_NO_3_ was applied per split application, resulting in a total of 2.314 g NH_4_NO_3_ (equivalent to 0.81 g N) for the N-high treatment.

All trials were conducted using the same setup to ensure consistency. The soil type used was CL ED73 Substrate. An alpha-lattice design was applied in each trial, with two replicates in GH1 and GH2 and three replicates in GH3. Ten seeds were sown in each 19 cm pot and maintained in a greenhouse at 20°C under a 12-hour light/dark cycle until the two-leaf stage. Seedlings were then transferred to a vernalization chamber at 4°C for 8 weeks. After vernalization, plants were moved to the PlantArray® platform (PA), where the temperature gradually increased from 10°C to 20°C over one week. During acclimatization, all plants were irrigated to maintain 70% of the water-holding capacity. On day 7, two irrigation treatments were applied: control (70% water capacity of the substrate) and drought stress (35% water capacity). The PA platform recorded pot weight and transpiration rate at three-minute intervals throughout the experiment.

### Plant growth conditions: field conditions

A field trial was conducted at the field station of Justus-Liebig-University Giessen in Groß-Gerau, Germany (49°56′27.5″ N, 8°30′06.8″ E). Fifty European winter wheat varieties were sown on 8 November 2022 at a density of 350 kernels m^−2^. The experiment was arranged as an alpha-lattice design with two irrigation treatments and two replicates per genotype and treatment combination. Crop management followed the station’s standard intensive practices, including herbicide and fungicide applications and fertilization with 120 kg N ha^−1^ (including N_min_). To impose post-anthesis drought stress in the respective treatment, irrigation was applied 206 and 232 days after sowing (∼6 and 32 days after heading) at a rate of 30 L m^−2^ each time. This resulted in a difference in total water supply of 60 L m^−2^ between irrigation treatments by harvest. Environmental data of the plant growth conditions were previously published.^85^

### Leaf image acquisition

A portable microscope (ProScope HR5, Bodelin Technology Co., Ltd, Oregon City, OR, USA) with a 400× magnification lens was used to image abaxial (AB) and adaxial (AD) leaf surfaces. Each image has a resolution of 2592 × 1944 pixels (width × height), corresponding to an image area of 0.806 mm^2^. In all environments and leaves measured, four technical replicates per leaf and per leaf surface were measured at four equidistant positions along the leaves (middle-base, leaf center, middle-tip, leaf tip). In the growth chamber, greenhouse, and field experiments, the third leaf, the third and sixth leaves, and the flag leaf were measured, respectively (Table 1). Biological replicates were two in the growth chamber experiments and greenhouse experiments GH1 and GH2. The greenhouse experiment GH3 and the field experiment had three and five biological replicates, respectively. In total, 26,893 images were taken. In the growth chamber experiments, stomatal images were taken in the third week after transplantation, when the third leaves were fully expanded. To increase the number of replicates, the measurements were repeated in the fourth and fifth weeks. One week after the onset of the irrigation treatments in the greenhouse experiments, images were taken from the fully expanded third leaves to assess the effects of soil drying on stomatal morphology. At this stage, the sixth leaves had not yet emerged. To evaluate the impact of drought on stomatal development, images of the sixth leaves were collected four weeks after the initiation of irrigation treatments, corresponding to the end of the tillering stage. In the field experiments, stomatal imaging was performed on flag leaves during the early grain-filling phase.

### Image processing annotations

Stomata were initially annotated manually using the graphical image annotation tool *roLabellm*.^86^ Each object was classified into one of five categories: complete, incomplete, blurry complete, blurry incomplete, or trichome structures. To establish representative ground-truth data for object detection, a total of 636 images were manually labelled. Two researchers labeled each stomata in these images, and if there were any discrepancies, two other researchers made the judgment together. This dataset incorporated the following: (i) both AB and AD leaf surfaces across all growth chamber conditions (HTHL, HTLL, LTHL, LTLL) as well as the open-field experiment; (ii) all genotypes; (iii) a high range of object densities; and (iv) a balanced representation of all stomatal classes.

### Model training

We give a brief overview of the parameters used for the model. The number of epochs was 1300, and the batch size was 16. During training, we adopted the AdamW optimizer, which updates model parameters based on both the current gradient and accumulated historical gradients, while explicitly applying weight regularization to improve generalization.^87^ The initial learning rate is set to 0.01 and gradually decreases following a cosine decay schedule to a final learning rate of 10^−4^, enabling rapid learning in the early stage and more refined parameter updates toward the end of training.

To avoid unstable training behavior at the beginning, a warm-up strategy was employed during the first 3 epochs, in which the learning rate and optimizer momentum were increased progressively from lower values. Specifically, the warm-up momentum was set to 0.8, and the bias parameters were trained with a slightly higher learning rate (0.1) to facilitate faster convergence of the detection head. The optimizer momentum, corresponding to the β1 parameter in AdamW, is set to 0.937, allowing the model to maintain consistent update directions and reducing oscillations during training.

To reduce overfitting and improve robustness, a weight decay of 0.0005 was applied as a regularization mechanism to discourage overly complex model parameters. In addition, a dropout rate of 0.3 was used to randomly deactivate a portion of neurons during training, encouraging the network to learn more generalizable representations. The loss function was designed to emphasize accurate object localization and classification, with a box loss gain of 7.5 and a classification loss gain of 0.5, while label smoothing is disabled to preserve sharp class boundaries.

Furthermore, various data augmentation techniques were applied to increase the diversity of training samples and improve the model’s generalization ability. The employed augmentation techniques included horizontal flipping, random cropping, hue and contrast adjustments, median blur, Gaussian blur, box blur, and motion blur.

### Stomatal and leaf trait calculations

Maximum stomatal conductance (*g_smax_*, mol m^−2^s^−1^) was estimated based on the following diffusion equation:^11,51^

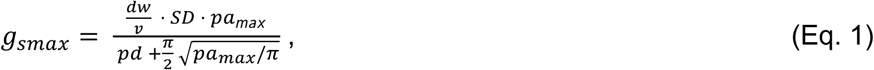

where *dw* is the diffusivity of water in air (2.49 × 10^−5^ m^−2^ s^−1^), and *v* is the molar volume of air (0.0224 m^3^ mol^−1^) at 25°C. Pore depth (*pd*, m) was assumed to be equal to stomatal width. The mean maximum stomatal pore area (*pa*_max_, m^2^) was estimated using the area formula for an ellipse, with the assumption that the major axis is the pore length (i.e., a quarter of the stomatal length) and the minor axis as half the pore length, *pd* is stomatal pore depth (m) was considered to be equal to the width of a fully turgid, inflated guard cell.^51,88^

For patterning statistics, the techniques and equations from a previous work were used.^15^ Stomatal evenness, *SE*_ve_, describes the regularity of stomatal distribution across the leaf surface with *N* stomata using a minimum spanning tree (MST) approach within a single image (Eq. 2):

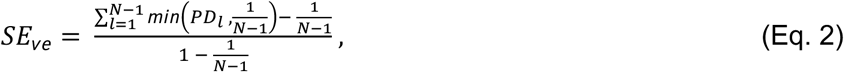

where *PD_I_* is the partial distance, defined as

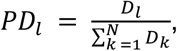

where *D_l_* is the Euclidean distance between stomata, and the stomata involved is branch *l* in the MST. *SD*_iv_ quantifies the divergence of stomatal positions by calculating the deviation from the center of gravity of *N* stomata:

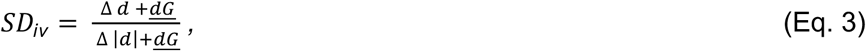

where △*d* is the sum of deviances from the center of gravity, △|*d*| is the absolute distance from the center of gravity, and *<u>dG</u>* represents the mean distance of the *N* stomata to the center of gravity. *SA*_gg_ represents the degree of stomatal clustering by calculating the ratio of the observed nearest-neighbor distance to the theoretical distance expected under a random distribution:

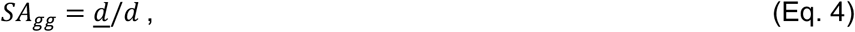

where *<u>d</u>* is the observed nearest-neighbor distance, and *d* is the theoretical nearest-neighbor distance considering an image’s stomatal density.

To quantify the proportion of phenotypic variance attributable to genetic differences among genotypes, broad-sense heritability (*H*^2^) was estimated as follows:^52^

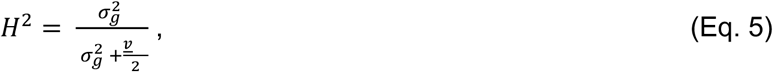

where 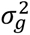 represents the genotypic variance and *v* is the mean variance of the differences between all possible pairs of genotypic best linear unbiased estimates (BLUEs) of derived genotype means.

### Linear mixed model formulation

To analyze the effect of certain environmental and plant variables (and their interactions) on each of the heritable stomatal traits, a linear mixed model (LMM) was created, and the Pearson correlation coefficients of the interactions were extracted (Fig. 5). Seven environmental and plant traits were analyzed, but not all of them were relevant for each growth condition (i.e. field, growth chambers, greenhouses). The following notation and their environmental conditions that it is relevant for are described as thus: genotype (G; field, GC, GH1, GH2, GH3), leaf side (LS, i.e. abaxial/adaxial; field, GC, GH1, GH2, GH3), temperature (T; GC, GH1, GH2, GH3), light (L; GC, GH1, GH2, GH3), leaf rank (LR; GH1, GH2, GH3), water treatment (W; GH1, GH2, GH3), and nitrogen treatment (N; GH3). Estimated marginal means (EMMs) were derived from the model using the *lme4* package in R (v.4.5.1)^89^, and the resulting correlations (Pearson correlation coefficients) of the individual traits (e.g., W, LS) and their interactions (e.g., N:LS, G:W:LR) were extracted and plotted (see Fig. 5).

### Calculation of the reverse genomic selection signal

To identify selection on the genomic basis, the reverse genomic selection approach described previously^56^ was applied, which relates estimated additive marker effects to allele-frequency changes over time. This method is specifically designed for highly quantitative and adaptive traits, where selection acts on many loci with small effects. The *G*^3^ statistic quantifies the genome-wide covariance between marker effects and allele frequency changes, which captures polygenic selection acting on many loci with small effects rather than being confined to strong single-locus signals.^56^ The reverse genomic selection analysis included all genotypes from the breeding panel for which both phenotypic and genotypic data were available (*n* = 49). One genotype (Asory) was excluded due to missing genotypic data. Allele frequencies were calculated based on the year of release for each genotype. To obtain robust estimates for allele frequency change and to reduce the influence of individual genotypes, the genotypes were grouped based on the release year. The oldest and most recent approximately 15% of genotypes were grouped into an early and a recent release-year group, respectively. Genotypes released between 1966 and 1975 (*n* = 8) formed the early group, and genotypes released between 2012 and 2016 (*n* = 7) formed the recent group. Allele frequency change for each SNP was then calculated as the difference between the early and recent genotype groups. Additive marker effects were estimated for each combination of trait, environment, and leaf side using the R package *rrBLUP*.^89^ The composite *G*^3^ statistic was subsequently computed with the R package *Ghat*,^90^ combining the allele-frequency changes with the estimated marker effects. Permutation testing with 5000 permutations was used to assess statistical significance.

