## Supplemental Data for "High-throughput stomatal phenotyping provides selection targets for stress-resilient wheat"

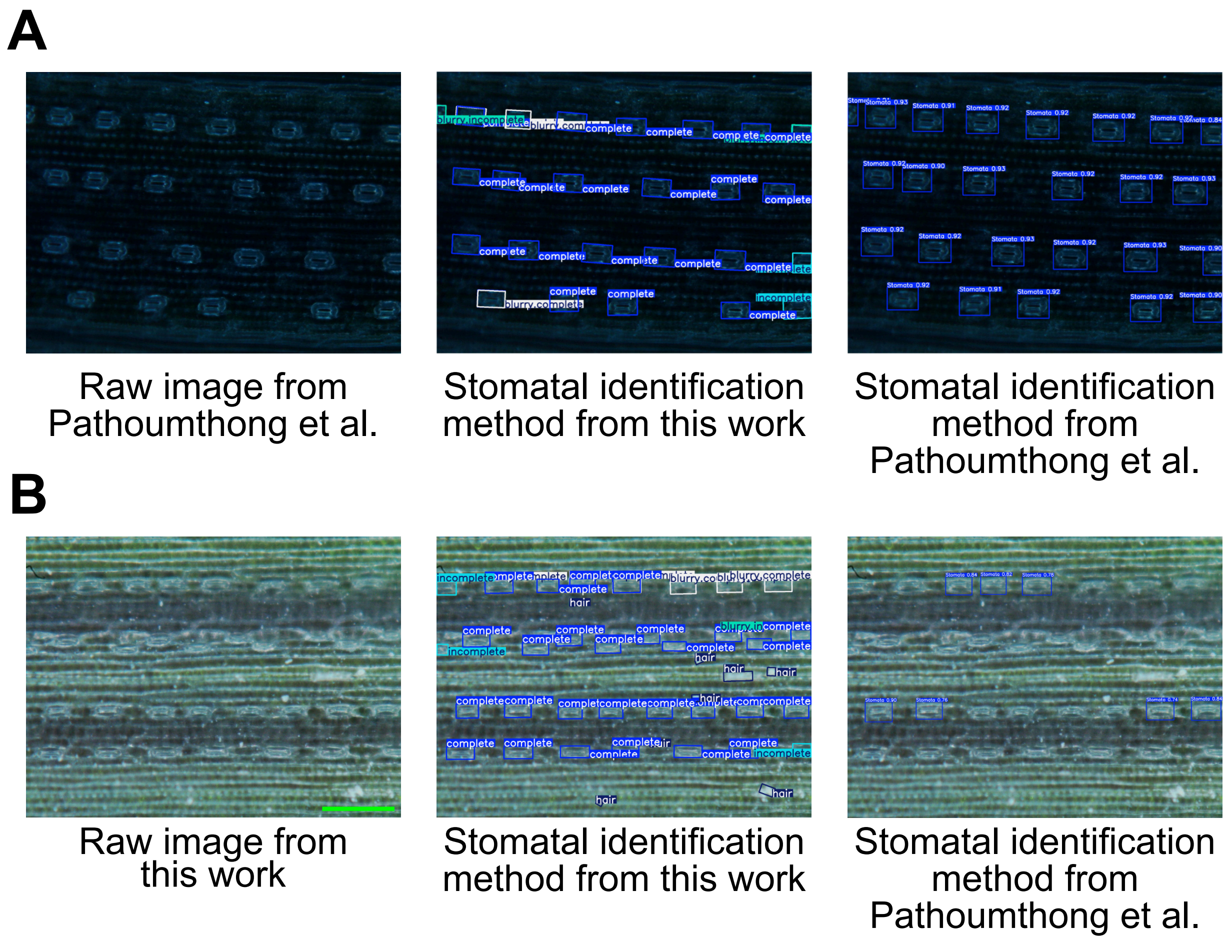


**Figure S1: The stomatal identification pipeline from our work accurately identifies stomata and hairs on leaves taken from different microscopes and leaves. Supplement to Figure 1.** Comparison of stomatal identification pipelines from this work and from Pathoumthong et al.[1] using **(A)** images from Pathoumthong et al. and **(B)** images from this work. The numbers on the Pathoumthong et al. images represent the probability that the detected object is a stomata. Scale bars = 200μm.

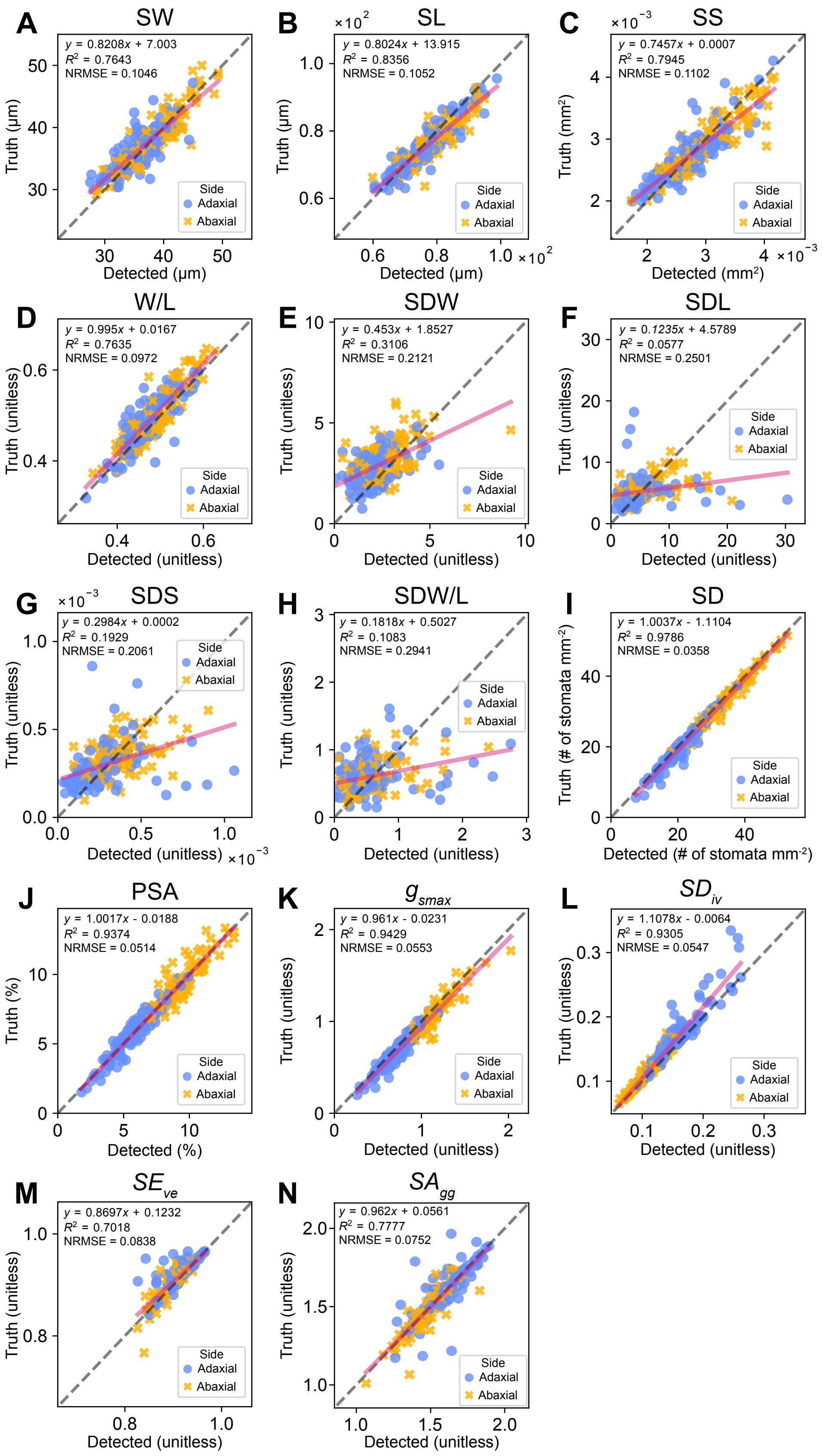
A SW
y = 0.8208x + 7.003
R2 = 0.7643
NRMSE = 0.1046
Truth (μm)
Detected (μm)
Side
Adaxial
Abaxial
B SL
y = 0.8024x + 13.915
R2 = 0.8356
NRMSE = 0.1052
C SS
y = 0.7457x + 0.0007
R2 = 0.7945
NRMSE = 0.1102
Truth (mm2)
Detected (mm2)
D W/L
y = 0.995x + 0.0167
R2 = 0.7635
NRMSE = 0.0972
E SDW
y = 0.453x + 1.8527
R2 = 0.3106
NRMSE = 0.2121
F SDL
y = 0.1235x + 4.5789
R2 = 0.0577
NRMSE = 0.2501
G SDS
y = 0.2984x + 0.0002
R2 = 0.1929
NRMSE = 0.2061
H SDW/L
y = 0.1818x + 0.5027
R2 = 0.1083
NRMSE = 0.2941
I SD
y = 1.0037x - 1.1104
R2 = 0.9786
NRMSE = 0.0358
Truth (# of stomata mm-2)
Detected (# of stomata mm-2)
J PSA
y = 1.0017x - 0.0188
R2 = 0.9374
NRMSE = 0.0514
Truth (%)
Detected (%)
K gsmax
y = 0.961x - 0.0231
R2 = 0.9429
NRMSE = 0.0553
L SDiv
y = 1.1078x - 0.0064
R2 = 0.9305
NRMSE = 0.0547
M SEve
y = 0.8697x + 0.1232
R2 = 0.7018
NRMSE = 0.0838
N SAgg
y = 0.962x + 0.0561
R2 = 0.7777
NRMSE = 0.0752
Truth (unitless)
Detected (unitless)

**<u>Figure S2: The stomatal identification and classification pipeline accurately calculates various stomatal morphological and patterning features. Supplement to Figure 1E.</u>**
Comparisons of various stomatal traits between the detected values by our pipeline (*x*-axis) and the ground truth values (*y*-axis) for the validation dataset (see Table 1). Each point represents either the mean value in each image or the value calculated from the entire image. The traits are as follows (see Table 2): **(A)** stomatal width (SW), **(B)** stomatal length (SL), **(C)** stomatal size (SS), **(D)** stomatal width-to-length ratio (W/L), **(E)** deviation in stomatal width (SDW), **(F)** deviation in stomatal length (SDL), **(G)** deviation in stomatal size (SDS), **(H)** deviation in stomatal width-to-length ratio (SDWL), (**I**) stomatal density (SD), **(J)** percentage of stomata area (PSA), **(K)** stomatal conductance ($g_{smax}$), **(L)** stomatal divergence ($SD_{iv}$), **(M)** stomatal evenness ($SE_{ve}$), and **(N)** stomatal aggregation ($SA_{gg}$). Blue circles represent values calculated for the adaxial side, whereas yellow squares represent values calculated for the abaxial side. The dashed line represents no difference between the ground truth and detected values. A pink line is drawn to represent a least-squares regression, the equation of which is in each subfigure, as well as the values of $R^2$ and the normalized RMSE (normalized by the mean).

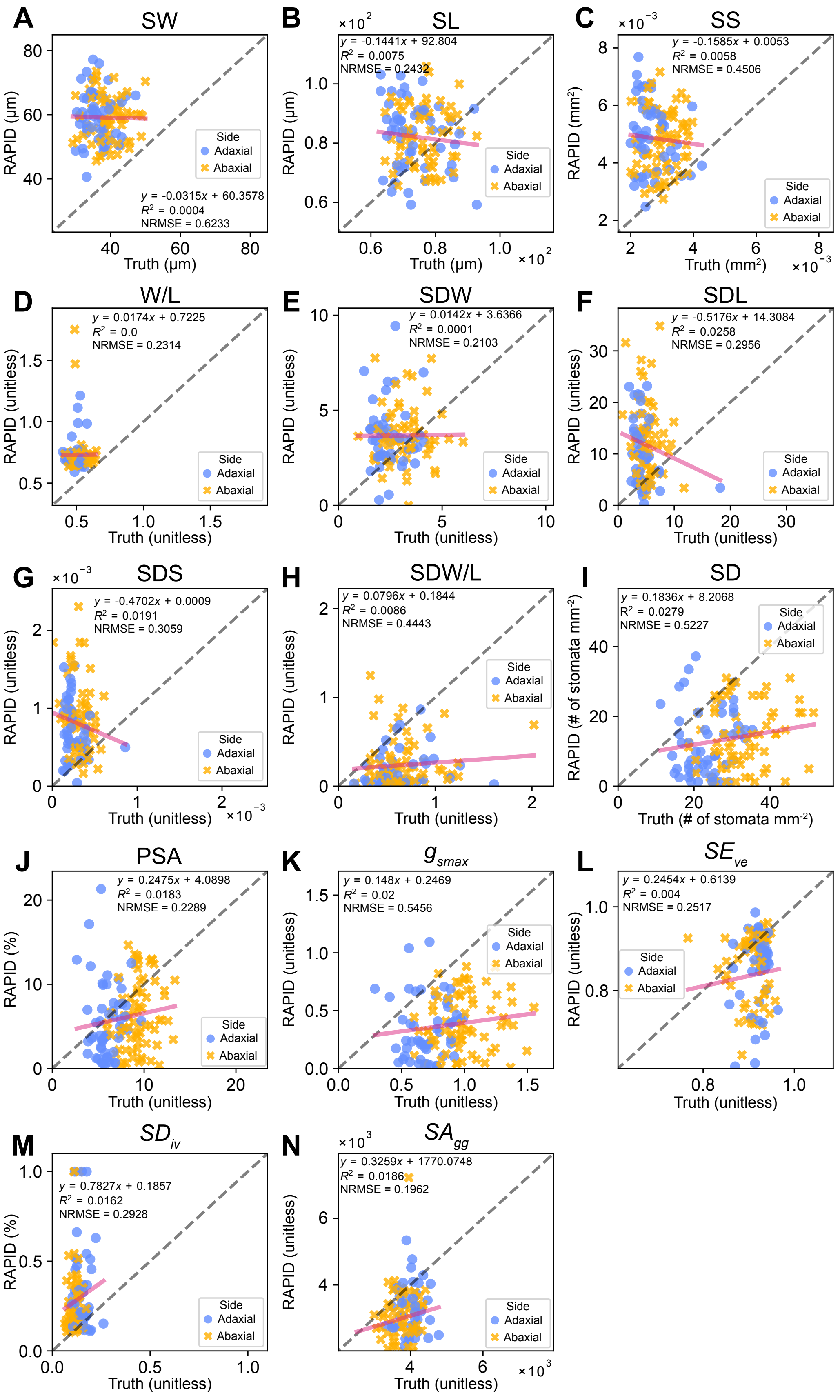
A
SW
RAPID (µm)
Truth (µm)
y = -0.0315x + 60.3578
R² = 0.0004
NRMSE = 0.6233
Side
Adaxial
Abaxial
B
SL
×10²
y = -0.1441x + 92.804
R² = 0.0075
NRMSE = 0.2432
RAPID (µm)
Truth (µm)
C
SS
×10⁻³
y = -0.1585x + 0.0053
R² = 0.0058
NRMSE = 0.4506
RAPID (mm²)
Truth (mm²)
D
W/L
y = 0.0174x + 0.7225
R² = 0.0
NRMSE = 0.2314
RAPID (unitless)
Truth (unitless)
E
SDW
y = 0.0142x + 3.6366
R² = 0.0001
NRMSE = 0.2103
F
SDL
y = -0.5176x + 14.3084
R² = 0.0258
NRMSE = 0.2956
G
SDS
y = -0.4702x + 0.0009
R² = 0.0191
NRMSE = 0.3059
H
SDW/L
y = 0.0796x + 0.1844
R² = 0.0086
NRMSE = 0.4443
I
SD
y = 0.1836x + 8.2068
R² = 0.0279
NRMSE = 0.5227
RAPID (# of stomata mm⁻²)
Truth (# of stomata mm⁻²)
J
PSA
y = 0.2475x + 4.0898
R² = 0.0183
NRMSE = 0.2289
RAPID (%)
K
gsmax
y = 0.148x + 0.2469
R² = 0.02
NRMSE = 0.5456
L
SEve
y = 0.2454x + 0.6139
R² = 0.004
NRMSE = 0.2517
M
SDiv
y = 0.7827x + 0.1857
R² = 0.0162
NRMSE = 0.2928
N
SAgg
×10³
y = 0.3259x + 1770.0748
R² = 0.0186
NRMSE = 0.1962

**Figure S3: The stomatal identification and classification pipeline of Pathoumthong et al. (2023) does not accurately quantify morphological and patterning traits in our images. Supplement to Figure S1.** Comparisons of various stomatal traits between the ground truth values (*x*-axis) and the pipeline of the Rapid method (Pathoumthong et al., 2023) (*y*-axis) for the validation dataset (see Table 1). Each point represents either the mean value in each image or the value calculated from the entire image. The traits are as follows (see Table 2): **(A)** stomatal width (SW), **(B)** stomatal length (SL), **(C)** stomatal size (SS), **(D)** stomatal width-to-length ratio (W/L), **(E)** deviation in stomatal width (SDW), **(F)** deviation in stomatal length (SDL), **(G)** deviation in stomatal size (SDS), **(H)** deviation in stomatal width-to-length ratio (SDWL), (**I**) stomatal density (SD), **(J)** percentage of stomata area (PSA), **(K)** stomatal conductance ($g_{smax}$), **(L)** stomatal evenness ($SE_{ve}$), **(M)** stomatal divergence ($SD_{iv}$), and **(N)** stomatal aggregation ($SA_{gg}$). Blue circles represent values calculated for the adaxial side, whereas yellow squares represent values calculated for the abaxial side. The dashed line represents no difference between the ground truth and detected values. A pink line is drawn to represent a least-squares regression, the equation of which is in each subfigure, as well as the values of $R^2$ and the normalized RMSE (NRMSE, normalized by the mean).

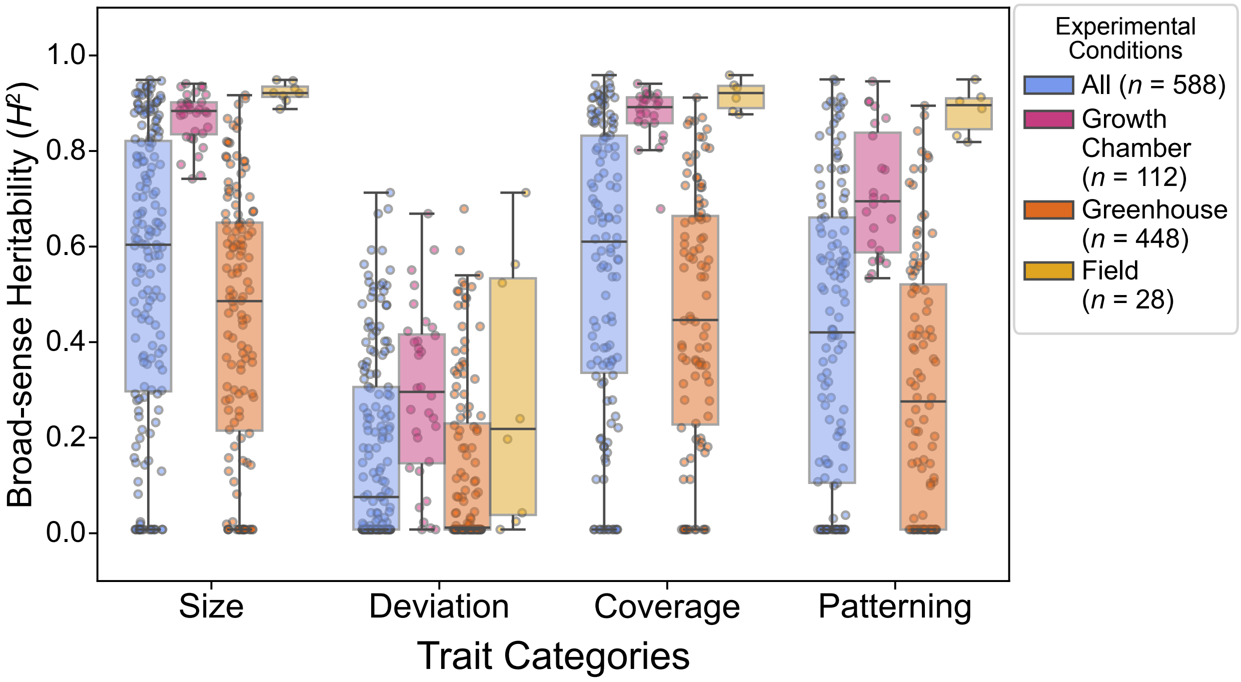


**Figure S4: Size, coverage, and patterning traits are highly heritable, particularly for growth chamber and field experiments. Supplement to Figure 2.** Broad-sense heritability of 14 stomatal traits grouped into four categories: size, deviation, coverage, and patterning. We assessed these trait categories in different environmental conditions (see Table 1 and Methods for more details): growth chambers (red dots and bars), greenhouses (orange dots and bars), and field conditions (yellow dots and bars). The blue bars and dots represent all 3 experimental conditions pooled together. The traits were aggregated as follows: Size: stomatal width (SW), stomatal length (SL), width-to-length ratio (W/L), and stomatal size (SS). Deviation: standard deviation of length (SDL), standard deviation of width (SDW), standard deviation of width-to-length ratio (SDWL), and standard deviation of size (SDS). Coverage: stomatal density (SD), percentage of stomatal area (PSA), and maximum stomatal conductance ($g_{smax}$). Patterning: stomatal evenness index ($SE_{ve}$), stomatal divergence index ($SD_{iv}$), and stomatal aggregation index ($SA_{gg}$).

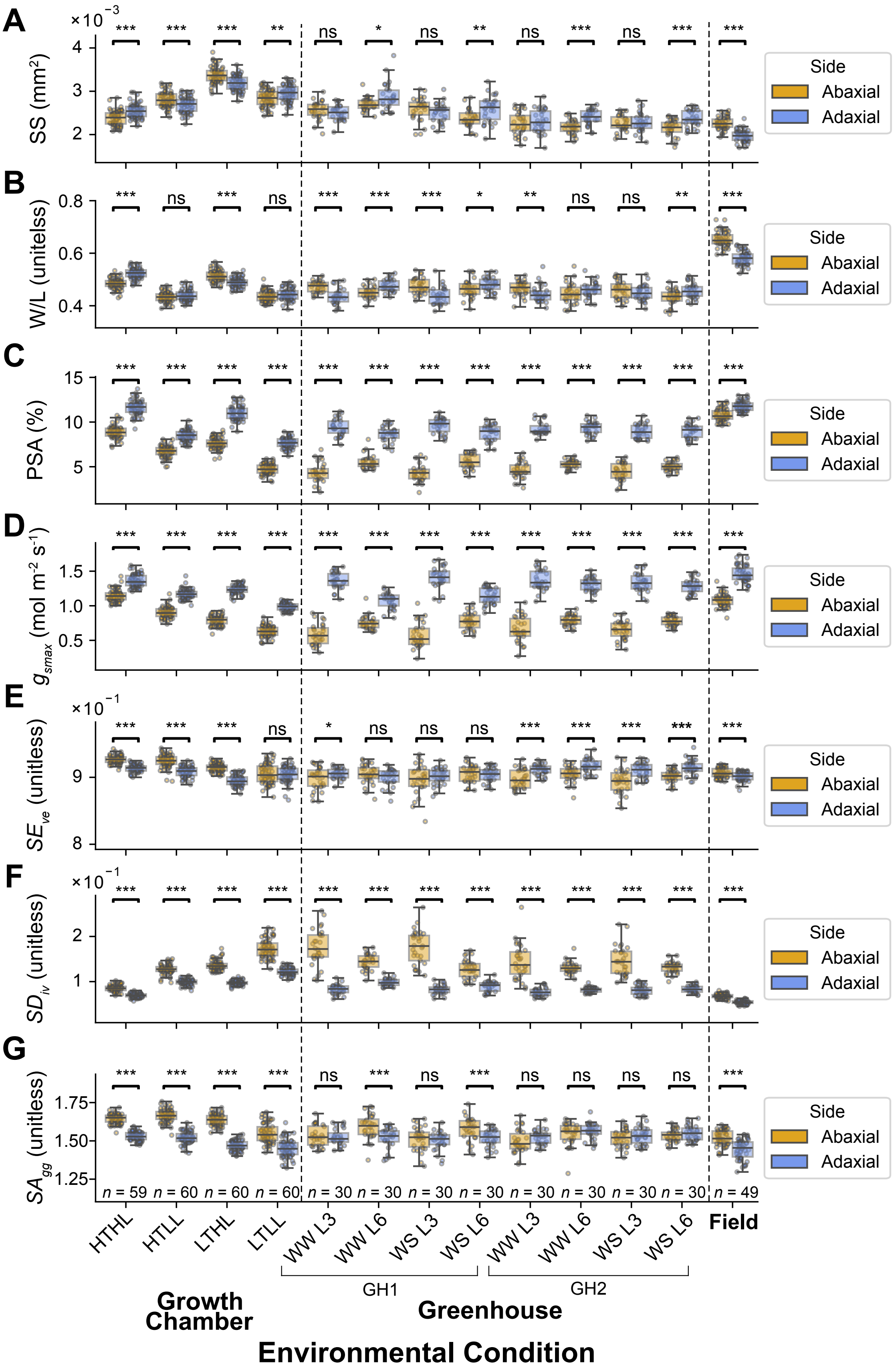
A
B
C
D
E
F
G
×10⁻³
×10⁻¹
SS (mm²)
W/L (unitelss)
PSA (%)
gsmax (mol m⁻² s⁻¹)
SEve (unitless)
SDiv (unitless)
SAgg (unitless)
Side
Abaxial
Adaxial
***
**
*
ns
n = 59
n = 60
n = 30
n = 49
HTHL
HTLL
LTHL
LTLL
WW L3
WW L6
WS L3
WS L6
Field
GH1
GH2
Growth Chamber
Greenhouse
Environmental Condition

**Figure S5: Analysis of stomata morphology highlights differences in genotypic and plastic responses depending on environmental conditions and leaf side. Supplement to Figure 3.** Comparison between **(A)** stomatal size (SS, $mm^2$), **(B)** stomatal width-to-length ratio (W/L, unitless), **(C)** percentage of stomatal area (PSA, %), **(D)** stomatal conductance ($g_{smax}$, mol $m^{-2}$ $s^{-1}$), (**E**) the stomatal evenness ($SE_{ve}$, unitless), (**F**) the stomatal divergence ($SD_{iv}$, unitless), and (**G**) the stomatal aggregation ($SA_{gg}$, unitless) of the abaxial (yellow) and adaxial (blue) sides of leaves in winter wheat cultivars grown under various environmental conditions: (1) four growth chamber treatments combining high temperature (HT) and low temperature (LT) with high light (HL) and low light (LL); (2) two greenhouse experiments (GH1 and GH2) that assessed the effects of drought stress (WS - water stressed; WW - well-watered) on the third leaf (L3) and sixth leaf (L6); and (3) a field experiment with no additional treatments. Each point represents the mean trait value for a single wheat cultivar in an environmental condition. A pairwise Student's *t*-test was conducted for each adaxial/abaxial pair with the following notation: * - $p < 0.05$, ** - $p < 0.01$, and *** - $p < 0.001$, and ns denotes non-significance.

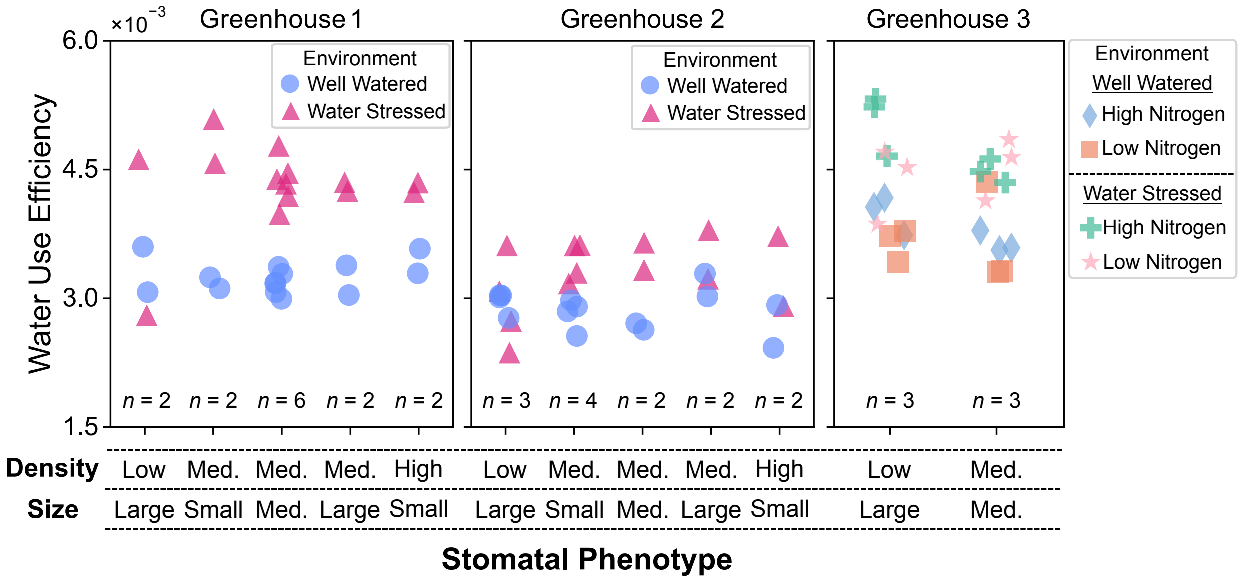


**Figure S6: Stomatal size and density do not influence water use efficiency across nitrogen and water stresses.** Water use efficiency (WUE, g $kg^{-1}$ $H_2O$) of winter wheat in different nitrogen and drought conditions in three greenhouses, stratified by stomatal morphology. Stomatal morphology was stratified into five groups: high stomatal density with large stomatal size, low stomatal density with large stomatal size, intermediate stomatal density with intermediate stomatal size, intermediate stomatal density with small stomatal size, and intermediate stomatal density with large stomatal size. Greenhouses 1 and 2 tested WUE between well-watered (blue circles) and water-stressed (pink triangles) conditions. Greenhouse three tested WUE between four conditions: well-watered and high nitrogen availability (blue diamonds), well-watered and low nitrogen availability (orange squares), water-stressed and high nitrogen availability (green crosses), and water-stressed and low nitrogen availability (pink stars). See Methods for how WUE was calculated.

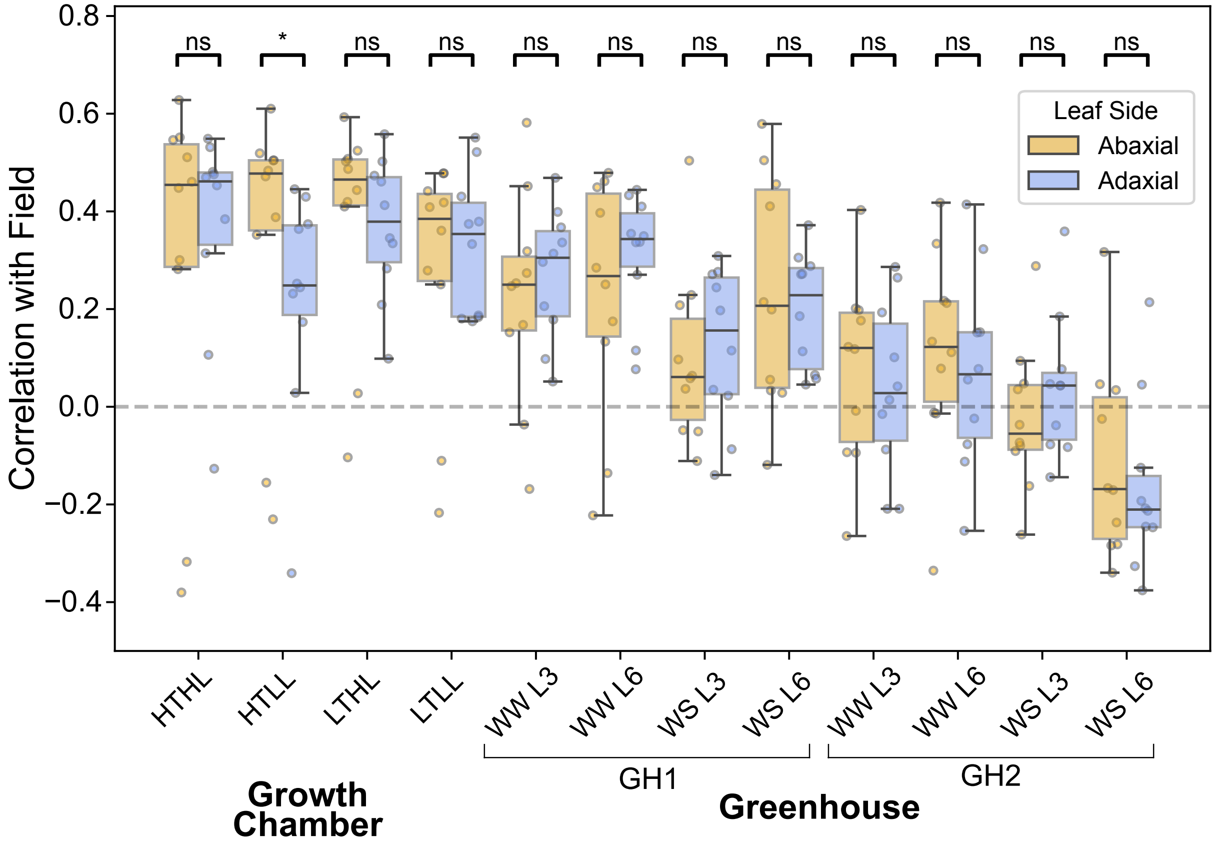


**Figure S7: The stomatal traits of third leaves grown in growth chambers correlate strongly with flag leaves grown in the field, regardless of leaf-side. Supplement to Figure 6.** Box plots of the Pearson correlation coefficients (*r*) of 10 heritable stomatal traits between field and controlled environments (greenhouse and growth chambers), stratified by leaf side (abaxial, yellow; adaxial, blue). See Fig. 4 for the traits used. The controlled environments include 1) four growth chamber treatments combining high temperature (HT) and low temperature (LT) with high light (HL) and low light (LL), and 2) two greenhouse experiments (GH1 and GH2) that assessed the effects of drought stress (WS - water-stressed; WW - well-watered) on the third and sixth leaves. Asterisks indicate significant differences between panels (*** - $p < 0.001$, ** - $p < 0.01$, * - $p < 0.05$), while ns denotes non-significance.

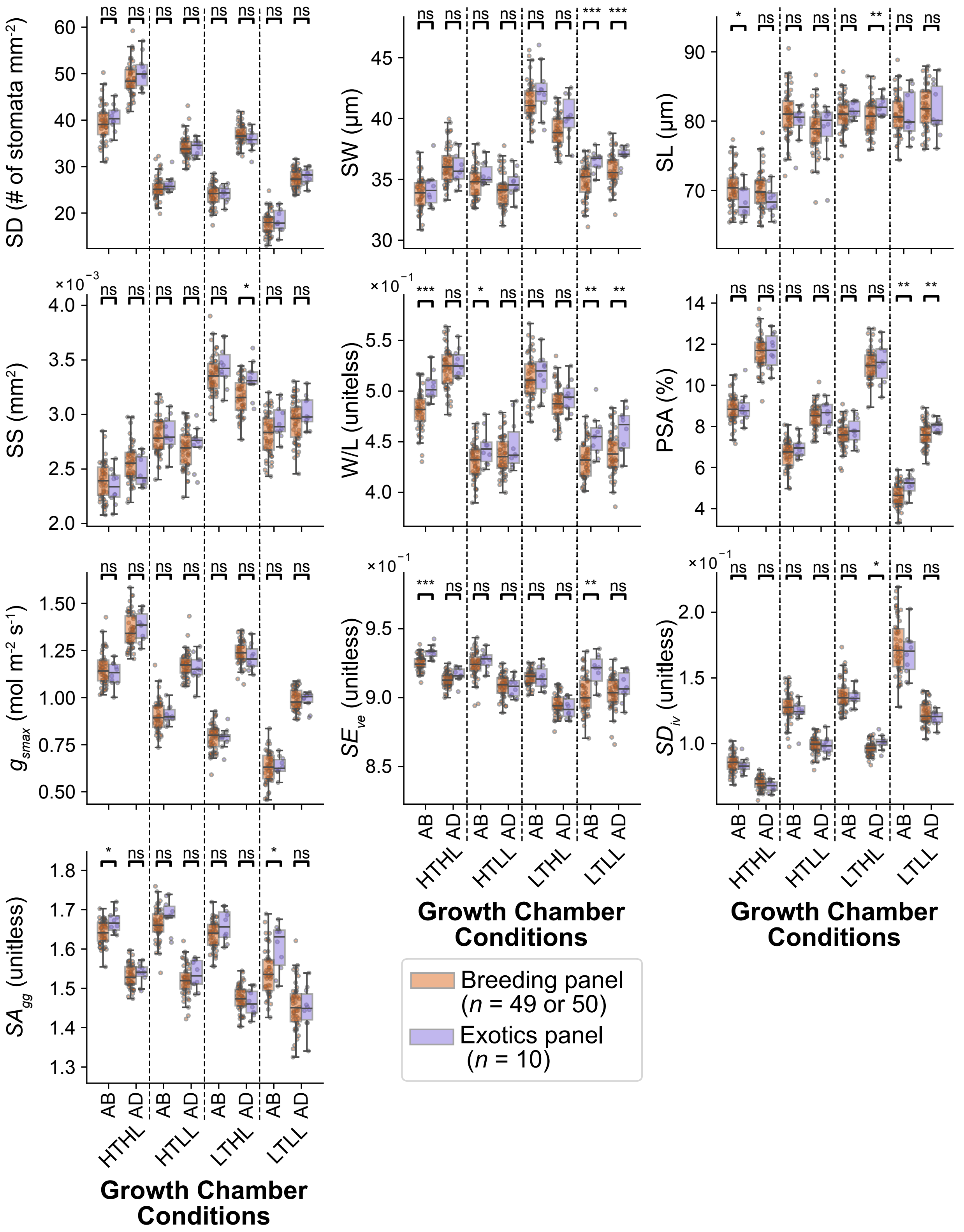


**Figure S8: German and non-German winter wheat varieties vary in their adaptation to low temperature and low light conditions.** Variation of stomatal traits between two panels: 50 winter wheat genotypes (breeding panel, red points) and 10 exotic cultivars (exotic panel, purple points and bars) grown under controlled environments. The traits are stomatal density (SD), stomatal width (SW), stomatal length (SL), stomatal size (SS), stomatal width-to-length ratio (W/L), percentage of stomata area (PSA), stomatal conductance ($g_{smax}$), stomatal evenness ($SE_{ve}$), stomatal divergence ($SD_{iv}$), and stomatal aggregation ($SA_{gg}$). Each point represents one cultivar. Controlled conditions included four growth chamber (GC) experiments combining two temperature levels (high temperature, HT; low temperature, LT) and two light levels (high light, HL; low light, LL). Measurements were taken from the third leaf. Asterisks indicate significant differences between panels (*** - $p < 0.001$, ** - $p < 0.01$, * - $p < 0.05$), while ns denotes non-significance.

| Cultivar | Breeder | Year of release | Grain quality | Country | Panel | Environment |
|---|---|---|---|---|---|---|
| Meister | RAGT/Mellinger | 2010 | A | DE | Breeding | Field, HTHL, HTLL, LTHL, LTLL, GH1 |
| KWS Santiago | KWS Lochow GmbH | 2011 | C | GB | Breeding | Field, HTHL, HTLL, LTHL, LTLL, GH1 |
| Paroli | DSV | 2004 | A | DE | Breeding | Field, HTHL, HTLL, LTHL, LTLL, GH2 |
| Patras | DSV | 2012 | A | DE | Breeding | Field, HTHL, HTLL, LTHL, LTLL, GH1, GH2, GH3 |
| Anapolis | NORDSAAT Saatzuchtgesellschaft | 2013 | C | DE | Breeding | HTHL, HTLL, LTHL, LTLL, GH2 |
| Biscay | Lochow-Petkus | 2000 | C | DE | Breeding | Field, HTHL, HTLL, LTHL, LTLL, GH1 |
| Cubus | Lochow-Petkus | 2002 | A | DE | Breeding | Field, HTHL, HTLL, LTHL, LTLL, GH1 |
| Forum | Nordsaat Saatzucht | 2012 | A | DE, EE, PL, SE | Breeding | Field, HTHL, HTLL, LTHL, LTLL, GH2 |
| Potenzial | DSV | 2006 | A | DE | Breeding | Field, HTHL, HTLL, LTHL, LTLL, GH1 |
| Gaucho | USDA-ARS, Oklahoma AES | 1993 | N/A | USA | Breeding | Field, HTHL, HTLL, LTHL, LTLL, GH2 |
| Tarso | Saatzucht Hadmersleben | 1992 | A | DE | Breeding | Field, HTHL, HTLL, LTHL, LTLL, GH1 |
| Hermann | Nickerson | 2004 | C | DE | Breeding | Field, HTHL, HTLL, LTHL, LTLL, GH1 |
| Tobak | W. von Borries-Eckendorf | 2011 | B | DE | Breeding | Field, HTHL, HTLL, LTHL, LTLL, GH1, GH2, GH3 |
| Pionier | DSV | 2013 | A | DE | Breeding | Field, HTHL, HTLL, LTHL, LTLL, GH2 |

| | | | | | | |
|---|---|---|---|---|---|---|
| Kalahari | Limagrain GmbH | 2010 | B | EU | Breeding | Field, HTHL, HTLL, LTHL, LTLL, GH2 |
| Intro | RAGT | 2011 | B | DE, FR | Breeding | Field, HTHL, HTLL, LTHL, LTLL, GH2 |
| Global | RAGT | 2009 | B | DE, AT | Breeding | Field, HTHL, HTLL, LTHL, LTLL, GH1, GH2, GH3 |
| Elixer | W. von Borris-Eckendorf | 2012 | C | DE | Breeding | Field, HTHL, HTLL, LTHL, LTLL, GH1 |
| Inspiration | Saatzucht Josef Breun GmbH & Co. | 2007 | B | DE | Breeding | Field, HTHL, HTLL, LTHL, LTLL, GH1, GH2, GH3 |
| Edgar | Limagrain GmbH | 2010 | B | DE | Breeding | Field, HTHL, HTLL, LTHL, LTLL, GH1 |
| Sponsor | Unisigma | 1994 | N/A | FR, IE | Breeding | Field, HTHL, HTLL, LTHL, LTLL, GH2 |
| Impression | Saatzucht Schweiger GbR | 2005 | A | DE | Breeding | Field, HTHL, HTLL, LTHL, LTLL, GH1, GH2, GH3 |
| Toronto | Strengs Erben | 1990 | A | DE | Breeding | Field, HTHL, HTLL, LTHL, LTLL, GH2 |
| Contra | Saatzucht Josef Breun | 1990 | C | DE | Breeding | Field, HTHL, HTLL, LTHL, LTLL, GH1 |
| Saturn | MPI | 1973 | B | DE | Breeding | Field, HTHL, HTLL, LTHL, LTLL, GH1 |
| JB Asano | Saatzucht Josef Breun GmbH & Co. | 2008 | A | DE | Breeding | Field, HTHL, HTLL, LTHL, LTLL, GH1 |
| Kerubino | Saatzucht Schmid Landau | 2004 | E | DE | Breeding | Field, HTHL, HTLL, LTHL, LTLL, GH1 |
| Orestis | Strube, Dr. H. | 1988 | B | DE | Breeding | Field, HTHL, HTLL, LTHL, LTLL, GH2 |
| Anthus | KWS Lochow GmbH | 2005 | B | DE | Breeding | Field, HTHL, HTLL, LTHL, LTLL, GH2 |

| | | | | | | |
|---|---|---|---|---|---|---|
| Herzog | Saatzucht Josef Breun | 1986 | A | DE | Breeding | Field, HTHL, HTLL, LTHL, LTLL, GH1, GH2, GH3 |
| Sorbas | Strube, Dr. H. | 1985 | B | DE | Breeding | Field, HTHL, HTLL, LTHL, LTLL, GH2 |
| Terrier | Nickerson | 2001 | B | DE | Breeding | Field, HTHL, HTLL, LTHL, LTLL, GH1, GH2, GH3 |
| Drifter | Nickerson | 1999 | B | DE | Breeding | Field, HTHL, HTLL, LTHL, LTLL, GH2 |
| Joss | Breustedt | 1972 | C | DE | Breeding | Field, HTHL, HTLL, LTHL, LTLL, GH1 |
| Disponent | Bayerische Saatzuchtgesellschaft | 1975 | A | DE | Breeding | Field, HTHL, HTLL, LTHL, LTLL, GH1, GH2, GH3 |
| Carisuper | Heidenreich und Eger | 1975 | A | DE | Breeding | Field, HTHL, HTLL, LTHL, LTLL, GH2 |
| Alidos | Saatzucht Hadmersleben | 1987 | E | DE | Breeding | Field, HTHL, HTLL, LTHL, LTLL, GH2 |
| Akratos | Strube, Dr. H. | 2004 | A | DE | Breeding | Field, HTHL, HTLL, LTHL, LTLL, GH2 |
| Diplomat | Firlbeck | 1966 | A | DE | Breeding | Field, HTHL, HTLL, LTHL, LTLL, GH1 |
| Benno | Bauer, G. | 1973 | E | DE | Breeding | Field, HTHL, HTLL, LTHL, LTLL, GH2 |
| Caribo | Heidenreich und Eger | 1968 | B | DE | Breeding | Field, HTHL, HTLL, LTHL, LTLL, GH2 |
| Cordiale | KWS UK Limited | 2003 | N/A | GB | Breeding | Field, HTHL, HTLL, LTHL, LTLL, GH2 |
| Premio | RAGT | 2006 | B | FR | Breeding | Field, HTHL, HTLL, LTHL, LTLL, GH1 |
| Camp Remy | Unisigma | 1980 | B | DE | Breeding | Field, HTHL, HTLL, LTHL, LTLL, GH1 |

| | | | | | | |
|---|---|---|---|---|---|---|
| Nimbus | Firlbeck | 1975 | B | DE | Breeding | Field, HTHL, HTLL, LTHL, LTLL, GH2 |
| Pegassos | Strube, Dr. H. | 1994 | A | EU | Breeding | Field, HTHL, HTLL, LTHL, LTLL, GH1, GH2, GH3 |
| Julius | KWS Lochow | 2008 | A | N/A | Breeding | Field, HTHL, HTLL, LTHL, LTLL, GH1 |
| RGT Reform | RAGT | 2014 | A | N/A | Breeding | Field, HTHL, HTLL, LTHL, LTLL, GH1 |
| Nordkap | Nordsaat | 2016 | A | N/A | Breeding | Field, HTHL, HTLL, LTHL, LTLL, GH1, GH2, GH3 |
| Asory | DANKO | 2018 | A | PL | Breeding | Field, HTLL, LTHL, LTLL, GH1 |
| Cajeme 71 | CIMMYT | 1971 | N/A | MEX | Exotics | HTHL, HTLL, LTHL, LTLL |
| BCD 1302/83 | Goertzen Seed Research | N/A | N/A | MD | Exotics | HTHL, HTLL, LTHL, LTLL |
| INTRO 615 | N/A | N/A | N/A | USA | Exotics | HTHL, HTLL, LTHL, LTLL |
| NS 22/92 | N/A | 1992 | N/A | SRB | Exotics | HTHL, HTLL, LTHL, LTLL |
| Labriego-Inia | INIA Chillan | 1981 | N/A | CHL | Exotics | HTHL, HTLL, LTHL, LTLL |
| Florida | Saatzucht Schweiger | 1985 | N/A | USA | Exotics | HTHL, HTLL, LTHL, LTLL |
| Hope | S. Dakota Agricultural Experiment Station | 1927 | N/A | USA | Exotics | HTHL, HTLL, LTHL, LTLL |
| Durin | Plant Breeding Int. Cambridge | 1986 | N/A | GN | Exotics | HTHL, HTLL, LTHL, LTLL |
| Mironovska 808 | Mironovska Research Institute | 1963 | N/A | UKR | Exotics | HTHL, HTLL, LTHL, LTLL |
| Mexico 3 | BAZ, Braunschweig Genetic Resources Centre | N/A | N/A | MEX | Exotics | HTHL, HTLL, LTHL, LTLL |

**Table S1:** Details of the cultivars examined in this study, encompassing the cultivar names, breeders, release years, grain quality classifications ‘E’, ‘A’, ‘B’, and ‘C’ (very high to very low baking quality) based on the German wheat quality standards, and their respective countries of origin. The breeder is classified according to the respective volume of the German Variety List (*Beschreibende Sortenliste*; BSL). The column “Environment” indicates which experiment the cultivars were tested on, including growth chamber conditions (HTHL - high temperature and high light; HTLL - high temperature and low light; LTHL - low temperature and high light; and LTLL - low temperature and low light), greenhouse conditions (GH1, GH2, and GH3), and field conditions. See Methods for specifics about the environmental conditions.

| Growing conditions | Experiment | Treatment | Investigated genotypes | Leaves screened | Images |
|---|---|---|---|---|---|
| Growth chamber | HTHL | Light and temperature | 59 | Third leaf | 5606 |
| | HTLL | Light and temperature | 60 | Third leaf | 2880 |
| | LTHL | Light and temperature | 60 | Third leaf | 5759 |
| | LTLL | Light and temperature | 60 | Third leaf | 2880 |
| | Validation | - | 60 | Third leaf | 240 |
| Greenhouse | GH1 | Drought stress | 30 | Third and sixth leaf | 1808 |
| | GH2 | Drought stress | 30 | Third and sixth leaf | 1920 |
| | GH3 | Drought stress and nitrogen | 10 | Third and sixth leaf | 1920 |
| Field | - | - | 49 | Flag leaf | 3880 |

**Table S2:** Summary of the experimental conditions of the three examined growing conditions. HTHL, HTLL, LTHL, and LTLL represent the temperature and light combinations in the growth chamber as follows: HTHL - high temperature and high light; HTLL - high temperature and low light; LTHL - low temperature and high light; and LTLL - low temperature and low light. In the HTHL condition, the cultivar “Asory” was not investigated due to seed availability. See Supplemental Table 1 for the list of cultivars in each growth condition.

| Dataset | Number of images | Environment | Leaf rank | Leaf side | Usage |
|---|---|---|---|---|---|
| *LTHL* | 120 | Growth chamber | Third leaf | Abaxial | Training |
| *LTHL* | 120 | Growth chamber | Third leaf | Adaxial | Training |
| *HTHL* | 50 | Growth chamber | Third leaf | Abaxial | Training |
| *HTHL* | 50 | Growth chamber | Third leaf | Adaxial | Training |
| *Gross-Gerau* | 28 | Field | Flag leaf | Abaxial | Training |
| *Gross-Gerau* | 28 | Field | Flag leaf | Adaxial | Training |
| *T22L600* | 120 | Growth chamber | Third leaf | Abaxial | Validation |
| *T22L600* | 120 | Growth chamber | Third leaf | Adaxial | Validation |

**Table S3: Summary of manually labelled images dataset**

| Parameter | Value | Description |
| --- | --- | --- |
| Epochs | 1300 | Number of iterations over the dataset |
| Batch size | 16 | Number of samples per gradient update |
| Optimizer | AdamW | see Henderson & Ferrari[2] |
| Initial learning rate ($lr_0$) | 0.01 | Controls the learning speed of the model |
| Final learning-rate fraction ($lr_f$) | 0.01 | Controls the reduction in the learning rate |
| Final learning rate | $1\times10^{-4}$ | $lr_0 \times lr_f$ |
| Momentum | 0.937 | Adam $\beta_1$ (see Henderson & Ferrari[2]) |
| Warm-up epochs | 3 | Prevents divergence during early training (see Kalra & Barkeshli[3]) |
| Warm-up momentum | 0.8 | See Kalra & Barkeshli[3] |
| Warm-up learning rate | 0.1 | See Kalra & Barkeshli[3] |
| Weight decay | $5\times10^{-4}$ | Prevents overfitting via regularization |
| Dropout rate | 0.3 | Prevents overfitting by dropping neurons |
| Box loss gain | 7.5 | See Xu et al.[4] |
| Classification loss gain | 0.5 | See Xu et al.[4] |
| Label smoothing | 0.0 | - |

**Table S4: Parameters that were used for training the stomatal identification and classification pipeline in our work.**
